# Brain transcriptomic profiling reveals common alterations across neurodegenerative and psychiatric disorders

**DOI:** 10.1101/2021.08.16.456345

**Authors:** Iman Sadeghi, Juan D. Gispert, Emilio Palumbo, Manuel Muñoz-Aguirre, Valentin Wucher, Valeria D’Argenio, Gabriel Santpere, Arcadi Navarro, Roderic Guigo, Natàlia Vilor-Tejedor

## Abstract

Neurodegenerative and neuropsychiatric disorders (ND-NPs) are multifactorial, polygenic and complex behavioral phenotypes caused by brain abnormalities. Large-scale collaborative efforts have tried to identify the genetic architecture of these conditions. However, specific and shared underlying molecular pathobiology of brain illnesses is not clear. Here, we examine transcriptome-wide characterization of eight conditions, using a total of 2,633 post-mortem brain samples from patients with Alzheimer’s disease (AD), Parkinson’s disease (PD), Progressive Supranuclear Palsy (PSP), Pathological Aging (PA), Autism Spectrum Disorder (ASD), Schizophrenia (Scz), Major Depressive Disorder (MDD), and Bipolar Disorder (BP)–in comparison with 2,078 brain samples from matched control subjects.

Similar transcriptome alterations were observed between NDs and NPs with the top correlations obtained between Scz-BP, ASD-PD, AD-PD, and Scz-ASD. Region-specific comparisons also revealed shared transcriptome alterations in frontal and temporal lobes across NPs and NDs. Co-expression network analysis identified coordinated dysregulations of cell-type-specific modules across NDs and NPs. This study provides a transcriptomic framework to understand the molecular alterations of NPs and NDs through their shared- and specific gene expression in the brain.

## INTRODUCTION

Neurodegenerative and neuropsychiatric disorders (ND-NPs) are multifactorial, polygenic, and complex behavioral phenotypes caused by changes in multiple underlying mechanisms^1,2^. In NDs, nerve cells become unable to respond to changes in their internal and external environments, eventually resulting in an impairment of brain function ^3–5^. At present, the most prevalent NDs^6,7^ are Alzheimer’s disease (AD), Parkinson’s disease (PD), progressive supranuclear palsy (PSP), as well as early preclinical manifestations of those such as pathological aging (PA). Because of the presence of amyloid plaques, but not tangles, and the absence of dementia, PA is considered to be either a prodrome of AD or a condition, in which there is resistance to the development of NFT and/or dementia^8^. Alteration of neuronal communications has been implicated in NDs, as well as in NPs such as schizophrenia (Scz), bipolar disorder (BP), autism spectrum disorder (ASD), and major depressive disorder (MDD), which are also among the major contributors to disability worldwide^9^. The etiology and mechanisms of NDs and NPs are elusive and a broad spectrum of causative genetic and environmental factors have been proposed ^10^.

Several investigations have shed light on the genetic heterogeneity within brain conditions and the degree of molecular similarities between closely related disorders^11–13^. Patterns of converging clinical and biological characteristics across NDs such as AD, PD, and PA have been lately discussed^14–16^. NPs have also shown symptomatic overlaps^17,18^. This demands uncovering condition-specific and overlapping pathological mechanisms across ND-NPs, which has been partly revealed by recent large-scale genome-wide association studies (GWAS)^19–21^. For instance, the genetic correlation between eight neuropsychiatric disorders revealed 3 different sub-groups with high levels of genetic overlap as well as multiple pleiotropic loci related to genes involved in neurodevelopment ^22^. Moreover, some effort has been made to relate shared genetic causes with shared transcriptomic alterations in the post-mortem brains in subsets of these disorders, revealing similar relationships among diseases even if not necessarily implicating the same genes ^21,23^.

Such integrative transcriptomic studies attempt to fill the functional gap and establish the degree of coupling between primary genetic causes and secondary events captured by the transcriptome in adult postmortem samples. However, a comprehensive characterization of gene expression changes in brain regions from individuals with major brain NDs and NPs compared with healthy subjects is missing. To address this shortcoming, we have uniformly analyzed a large collection of bulk RNA-Seq samples from post-mortem brain regions of subjects with NDs and NPs produced by different studies. The results of our meta-analysis revealed similarities of transcriptomic alterations between NDs and NPs which have also been observed in brain regions such as frontal and temporal lobes across the conditions. We additionally found coordinated downregulation of neuron-specific co-expression modules in both NDs and NPs, while oligodendrocyte and astrocyte modules showed mainly upregulation across conditions.

## RESULTS

### Samples characteristics and clustering

We analyzed 4,711 RNA-Seq samples produced by 19 different labs^8,21,24–41^ from patients with AD (n = 906 samples), PD (n = 29), PA (n = 58), PSP (n = 168), Scz (n = 535), ASD (n = 187), MDD (n = 240), BP (n = 510), and non redundant matched controls (n = 2,078) pooled across all studies, obtained from seven major brain regions (**Fig. 1 & S1**, for further details see **Table S1 & S2**). To produce a uniform gene expression quantification that could be compared across different datasets, the samples were processed using the Grape RNA-Seq pipeline^42^ and underwent normalization and quality control (see **Methods & Fig. S2-S9**). t-Distributed Stochastic Neighbor Embedding (tSNE)^43^ analysis in combination with principal component analysis (PCA) was used to produce a latent representation of the samples based on their gene expression profiles. Globally, NDs and NPs clustered separately, driven by the disjoint clustering of AD and Scz, respectively (**Fig. 2a**). PA and PSP clustered together very distinctively, but this could be confounded because samples for these conditions are coming from exactly the same regions. ASD clustered into two different groups, likely reflecting the brain region from which samples originated. This could indicate that ASD may correspond to different molecular conditions. Control samples showed little structure, supporting the effectiveness of our normalization, whereas, condition samples demonstrate a stronger structure (**Fig. S10**). To some extent, this also happens with MDD. Finally, BP shows partly clustering with Scz based on the origin of the regions (**Fig. 2a**).

**Figure 1.**
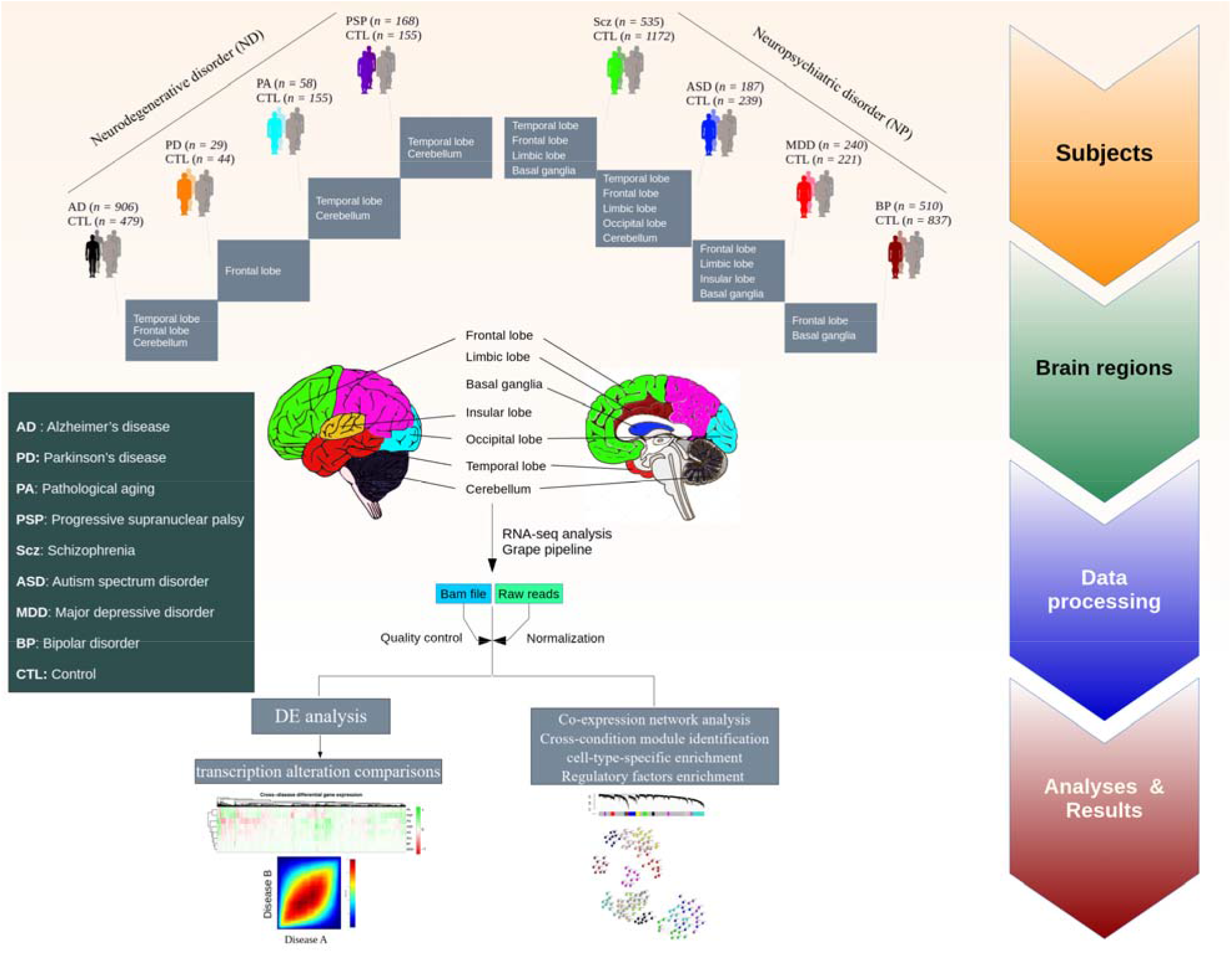
Schematic of the study design and samples used for gene expression analysis via an RNA-Seq pipeline. Post-mortem brain RNA-Seq data were obtained from subjects with AD (n = 906 samples), PD (n = 29), PA (n = 58), PSP (n = 168), Scz (n = 535), ASD (n = 187), MDD (n = 240), BP (n = 510), and matched controls (n = 2078) (see **Supplementary Table S1&S2** and **Fig. S1**).

**Figure 2.**
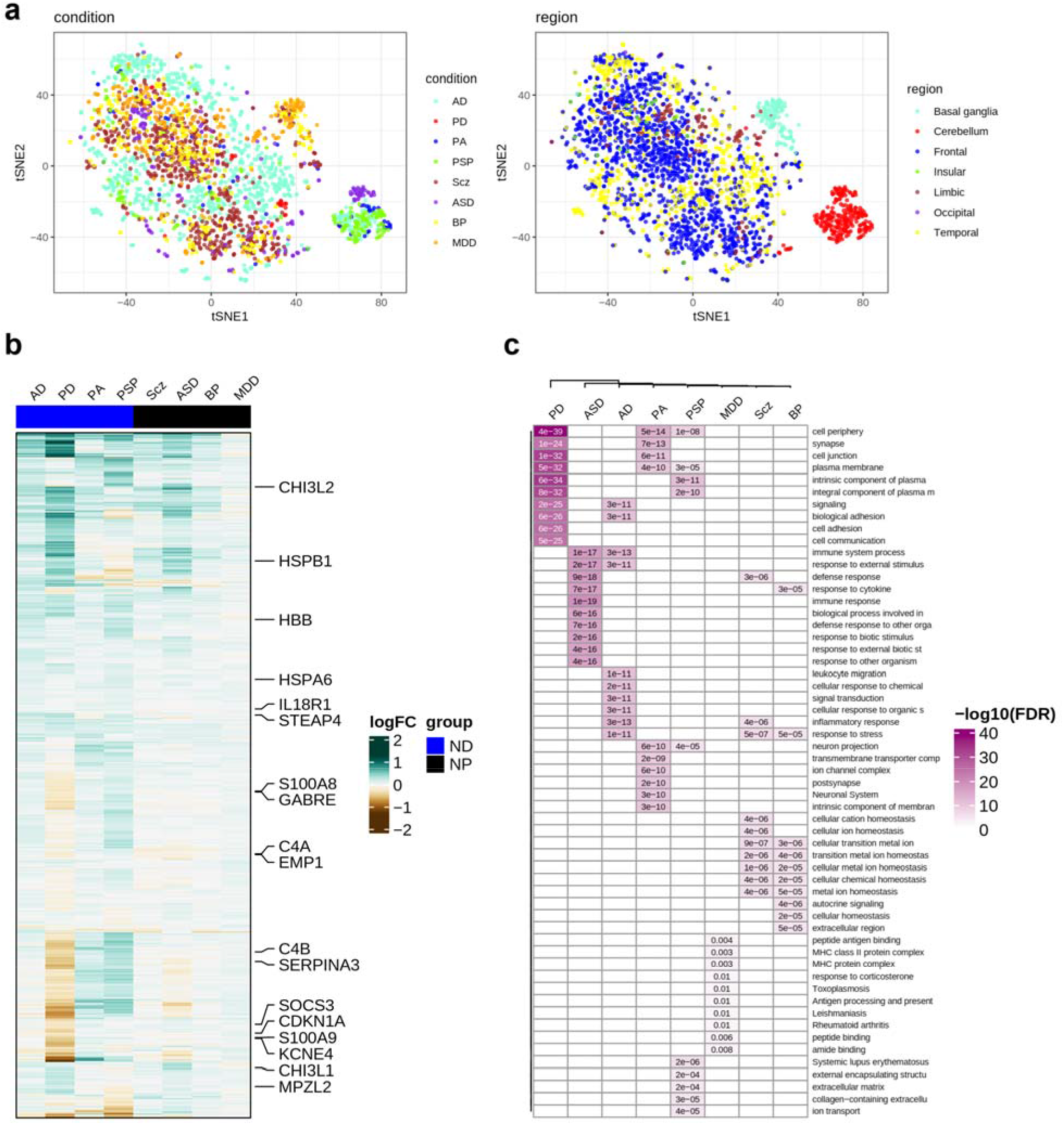
Condition-specific transcriptome alterations. (**a**) tSNE visualization of the pooled samples as colored by conditions (left) and brain regions (right). (**b**) A heatmap of differentially expressed genes across neurodegenerative disorders (ND) and neuropsychiatric disorders (NP). The row labels represent the genes differentially expressed in at least 5 conditions. (P < 0.05 & |log2FC| > 0.5). (**c**) Conditions-specific gene enrichment analysis. Top significantly enriched pathways are represented for significantly differentially expressed genes across conditions (FDR-corrected p-value < 0.05).

### Condition-specific differential gene expression (DGE)

Condition-specific differential gene expression (DGE) analyses were performed using a linear mixed-effect model (see **Methods**). These analyses provided insights regarding transcriptional changes for the pathobiology of each condition (**Supplementary Data 1**). In total, we found 4419 genes differentially expressed in at least one condition (**Fig. S11a,b)**. Significant overlap between condition-specific DEGs and those obtained from individual datasets represented the reproducibility of the results (**Fig.S12**). Most DEGs were exclusive of one single condition, and we did not find a single gene shared across all the conditions, but there were genes shared across a number of conditions **(Fig. 2b & Fig. S13**). For instance, *CHI3L1* (which codes for a protein also known as YKL-40) and *CHI3L2* that have been associated with astrocytic reactivity and neuronal damage^44,45^, are deregulated in all conditions except in MDD. Gene enrichment analysis performed using DEGs for each condition showed that most functional alterations were quite condition-specific, but functions related to and to the immune response were shared among some conditions (FDR-corrected p-value < 0.05; **Fig. 2c**).

### Cross-condition transcriptome overlap observed across NDs and NPs

To investigate the similarity between transcriptome alterations underlying the NDs and NPs, we compared the genes’ fold change (FC) in expression in cases vs controls between conditions, by explicitly computing the pairwise correlation of logFCs from 15,819 shared genes (See **Methods, Supplementary Fig. S14a** and **Supplementary Data 2**). This set of genes showed significant overlap with the list of common genes across psychiatric disorders from Gandal et al^21^. study (odd ratio = 4.5, FDR-corrected p-value < 0.001). Scz and BP represented the strongest correlation with the overlap of both downregulated and upregulated genes (**Fig. 3a & Fig. S14b**). According to the transcriptional alterations, ASD clustered together with NDs, rather than within the other NPs (**Fig. 3b**). Within NP’s, Scz and BP clustered closer, and within NDs, PA and PSP. MDD clustered separately from the rest of the conditions (**Fig. 3b**). These results were confirmed by computing correlations of the logFCs from the same set of genes across individual datasets to check for reproducibility (**Fig. S14c**).

**Figure 3.**
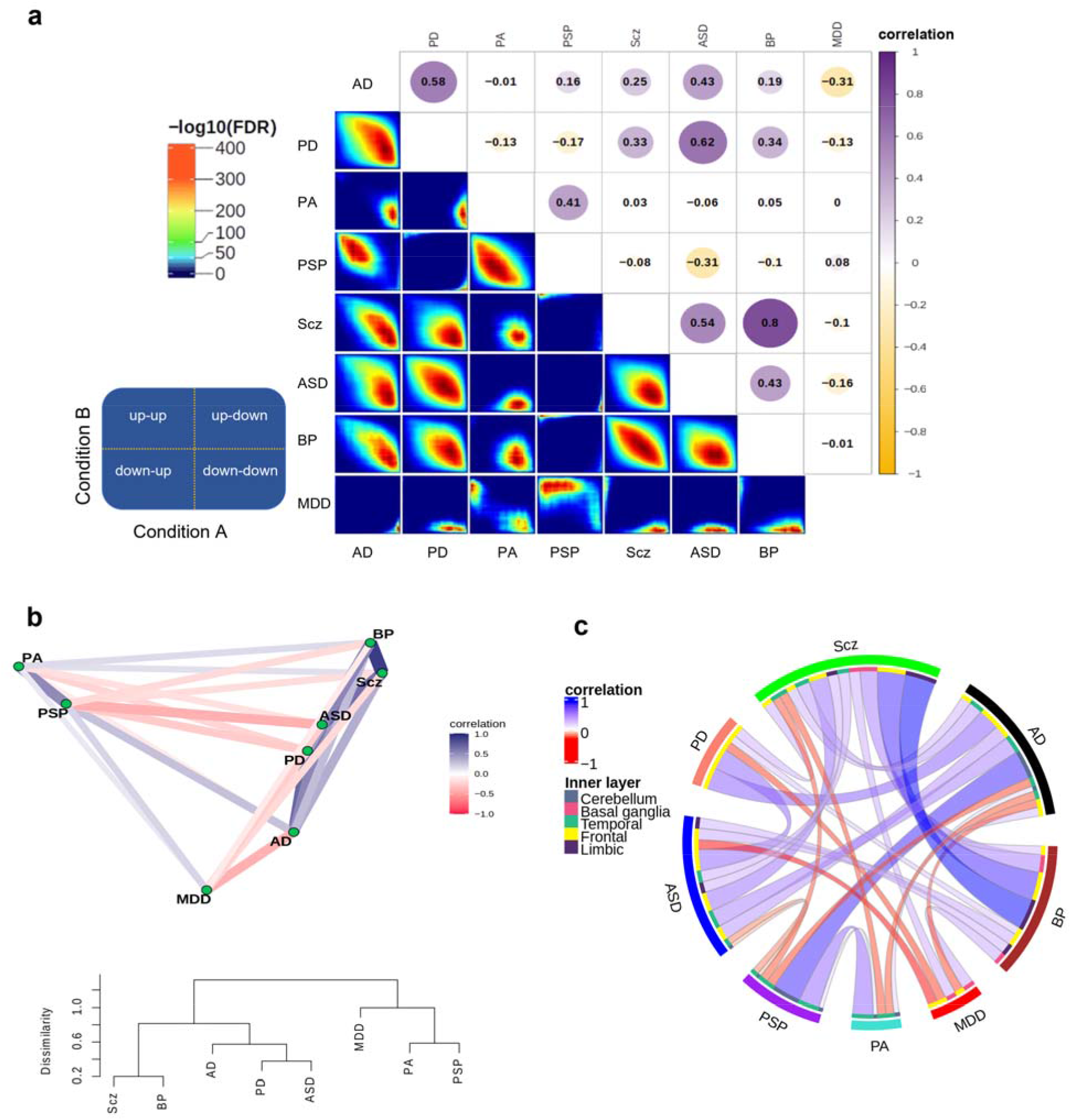
Similarity of transcriptional alterations across conditions. (**a**) Correlation plot (**top**) shows transcriptome alterations overlap obtained by computing Spearman’s correlations using logFC values of the shared genes between the conditions. Rank-rank hypergeometric overlap (RRHO; **bottom**) depicts direction (upregulation and downregulation) of the logFC overlaps. The guide panel represents the cross-condition overlapping relationship. Signals in the upper left quadrant display an overlap for shared upregulated genes, while those in the bottom right quadrant depict shared downregulated genes. The color bar displays the degree of significance of the overlap (Fisher’s exact test with FDR < 0.05). (**b**) Correlation network (**top**) and a tree dendrogram (**bottom**) obtained from pairwise correlations corresponding to **a**., show relationship of the conditions based on transcriptome alterations. (**c**) A circos plot demonstrating correlations of transcriptional alterations across conditions. Only significant correlations after FDR correction (FDR < 0.05) with a cut-off of absolute correlation > 0.1 are displayed here (see **Supplementary Fig. S17**). The outer layer represents conditions, while the inner layer displays brain regions defined by colors.

### Region-specific differential gene expression

Although some regions were present for only a limited number of conditions, brain-region-specific DGE revealed a set of overlapping genes that were frequently differentially expressed in multiple brain regions across conditions or vice versa (**Fig. S15 & Supplementary Data 3**). These genes included *SERPINA3* associated with neuropathies and the formation of amyloid-fibrils in condition^46–48^, *S100A8/A9, Socs3, ILR1L1* associated with inflammation^49,50^, *APOL4* involved in lipid metabolism^51^, and *NPAS4* which regulates the excitatory-inhibitory balance within neural circuits^52^. We built classifier models using the expression of DEGs from temporal and frontal regions (the most present regions across conditions) to explore the discrimination power of transcriptomic profiles between condition and control samples. High prediction accuracy was obtained for PD (78%) and AD (79%) in the frontal and temporal cortex, respectively (**Fig. S16**). Analysis of the similarities in transcriptional alterations between different conditions in the individual brain regions separately generally recapitulated the findings in **Fig. 3a**. BP and Scz showed a high positive correlation in all the regions in which these conditions were assayed (**Fig. 3c & Fig. S17**). MDD showed little correlation with the other conditions across different regions. Some conditions, however, showed similar or distinct transcriptome alterations depending on the region. Thus, transcriptomic changes underlying PSP and AD were highly correlated in the cerebellum, but negatively associated in the temporal lobe (**Fig. 3c & Fig. S17**). In some cases, therefore, apparently similar phenotypic outcomes are the consequence of different molecular events in different brain regions.

### Network analyses identified condition-specific and shared transcriptional signatures

To connect molecular alterations with the phenotypes of NDs and NPs, via their impact on the cellular composition of the brain, we constructed co-expression networks over all combined normalized datasets for the same set of shared genes using weighted gene co-expression network analysis (rWGCNA)^53^ and obtained seventeen co-expression modules (**Fig. 4a & Supplementary Data 4**). Module M0 comprises the genes that are not included in specific modules. We assigned cell types to each module based on the overlap between the genes in the module and the genes defining the cell type as in the PanglaoDB database^54^. A number of modules were enriched for cells of a specific type: M2 for oligodendrocytes and Schwann cells, M5 for astrocytes and Bergmann glia, M10 for microglia, M8 and M13 for neurons, and M14 for ependymal cells (**Fig. 4b**). These modules were actually the most distinct from the rest according to the correlation analysis and MDS (**Fig. S18**). The assignments were confirmed using an independent single-cell expression dataset^55^ composed of five main brain cell types including neurons, astrocytes, oligodendrocytes, microglia, and endothelial cells (**Fig. S19a**). We also identified the hub genes (the genes with the highest intramodular connectivity for each module (**Fig. S19b**). The functional categories enriched in cell-type-specific modules were broadly consistent with the cell type assignments (**Fig. 4c & Supplementary Data 5**).

**Figure 4.**
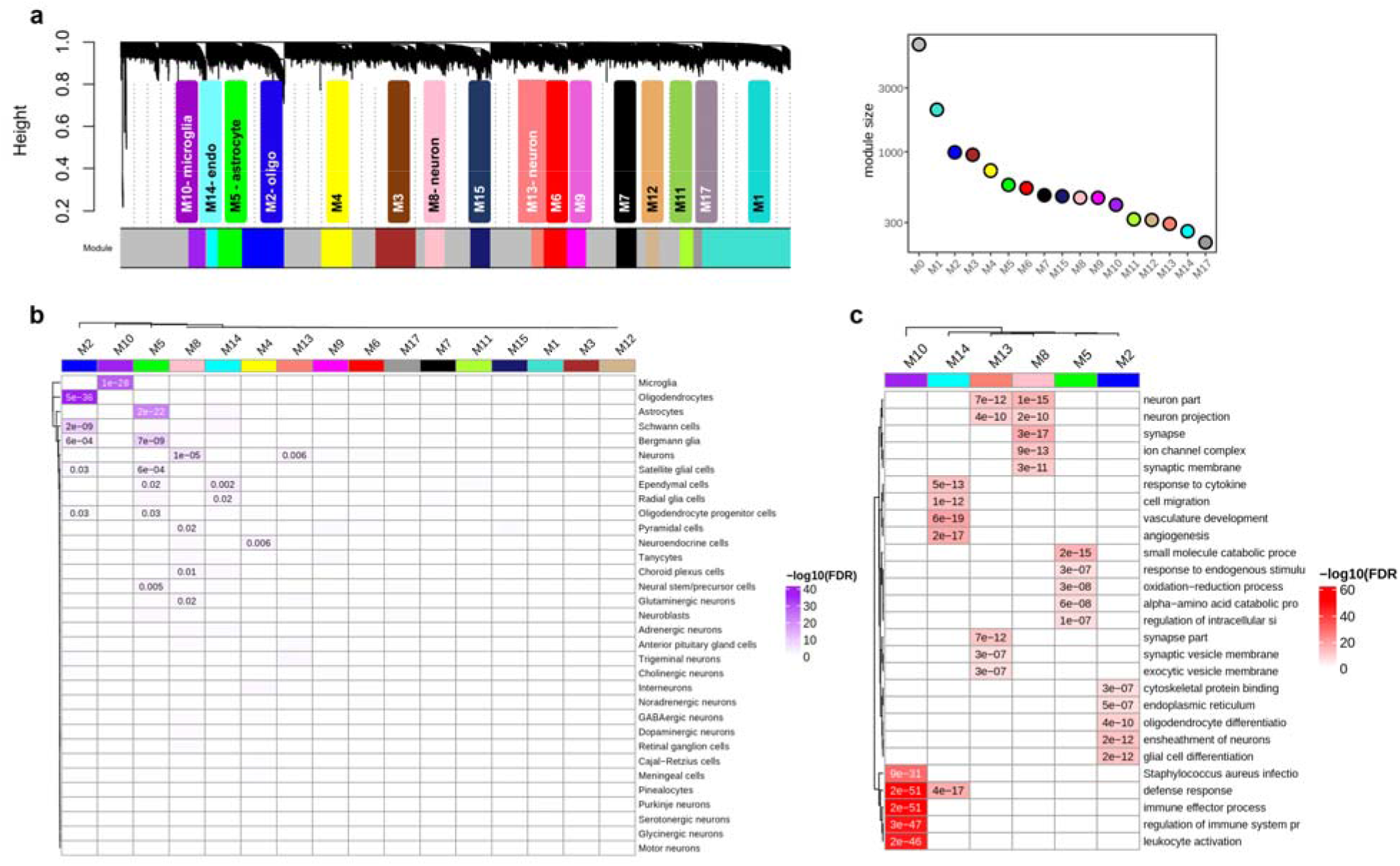
Cross-condition co-expression modules identified by network analysis. **(a)** A dendrogram plot displaying co-expression modules obtained from topological overlap of 15,819 shared genes between conditions. Each color represents an individual module and the grey color (M0) contains genes that are not included into a specific module. The corresponding plot at the right shows the number of genes within each module. (**b**) Enrichment of co-expression modules for brain cell types, measured by comparing genes within each module to the brain single-cell dataset from PanglaoDB^54^ (see also **Supplementary Fig. S19a**). **(c)** Heatmap plot of gene ontology enrichment for cell-type-specific modules using top five significant pathways for each module. Color key shows -log_10_(FDR).

We next identified the differential expression of cell-type-specific modules in each condition from the expression of eigengenes (the first principal component of the expression matrix of the corresponding module, as a representative of an entire co-expression module) in each module using a linear mixed model (see **Methods, Fig. 5a & Supplementary Data 6**). In addition, the differential expression of top hub genes within these modules was measured for each condition (**Fig. 5b**). Then, we could relate NDs and NPs to cellular alterations in the brain.

**Figure 5.**
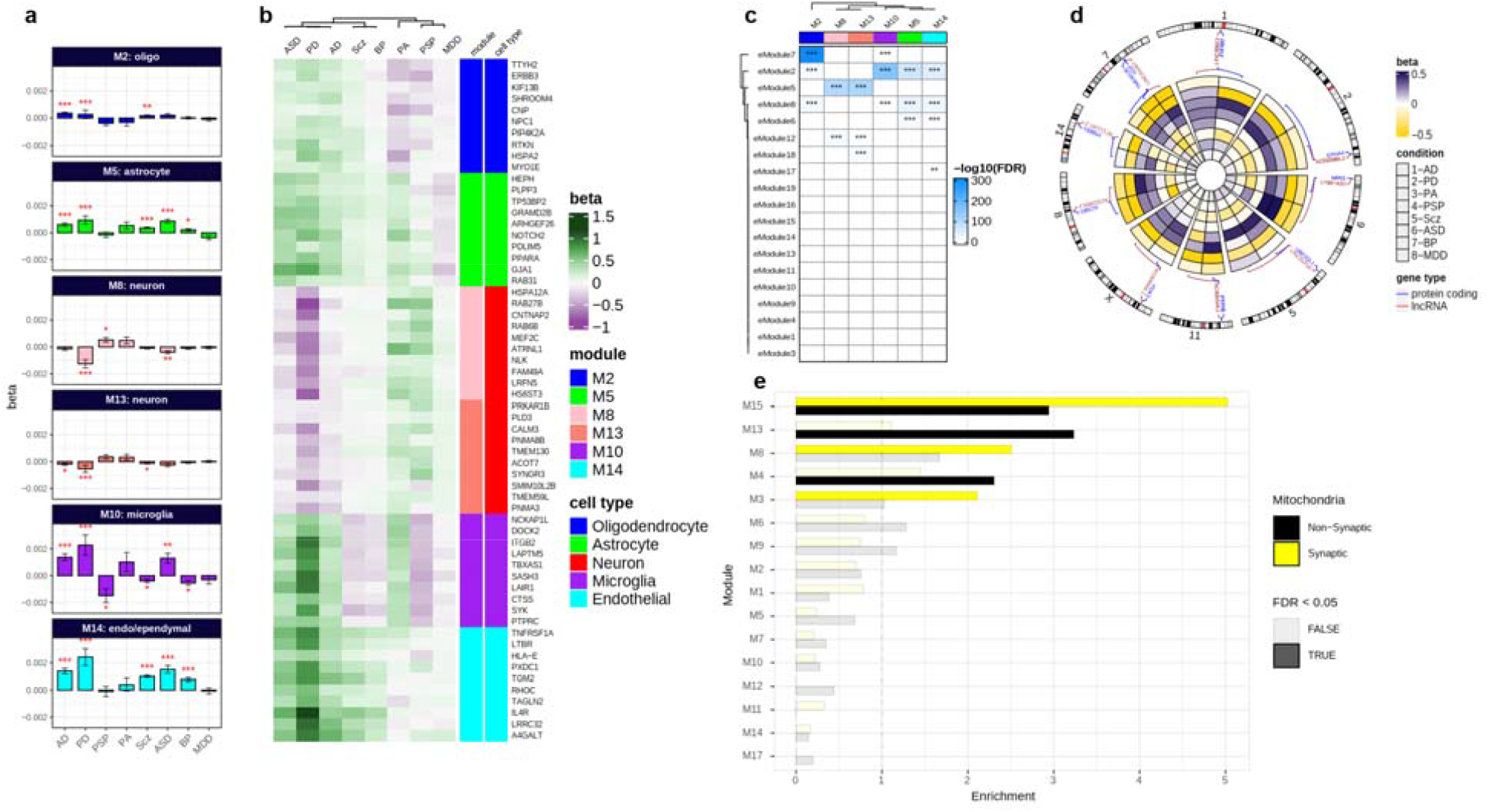
Co-expression gene modules characterizations. (**a**) Differential expression of cell-type-specific modules across conditions. □ values on y-axis computed by linear mixed effect model show relationship of modules eigengenes with conditions. Bar plots show mean ± SE values (**b**) Differential expression of top hub genes within cell-type-specific modules across conditions. Brain cell-type-specific modules are annotated with colors. (**c**) Enrichment of brain enhancer RNAs for cell-type-specific modules. The overlap between co-expression modules and eRNA modules from an independent dataset^65^ was computed by Fisher’s exact test (FDR < 0.05). Color key shows the -log10 (FDR-corrected p-values; *FDR < 0.05, **FDR < 0.01, ***FDR < 0.001; see also **Supplementary Fig. S20**). (d) A circular heatmap showing expression of protein coding and their flanking lncRNAs in neuron modules M8 and M13 across conditions. Conditions from outer to inner represent AD to MDD. Gene types are colored by blue (protein coding) and red (lncRNA) (See also **Supplementary Fig. S21**). (**e**) The enrichment of co-expression modules for mitochondrial transcriptomes. An independent study that previously reported synaptic and nonsynaptic mitochondria co-expression modules was obtained and compared to co-expression modules in this study using Fisher’s exact tests (FDR < 0.05). Yellow and black colors represent enrichment of synaptic and nonsynaptic mitochondrial transcriptomes for co-expression modules.

Thus, neuron-specific modules (M8 and M13) were broadly downregulated across AD,PD,ASD,Scz,and BP, but upregulated in PA and PSP. The oligodendrocyte module M2 was upregulated in all conditions except PA and PSP (**Fig. 5a,b**), and consistently enriched with genes involved in oligodendrocyte and glial cell differentiation (**Fig. 4c**). The microglia-associated module (M10) which was upregulated in neurodegenerative diseases AD and PD, PA and a psychiatric disorder ASD represented enrichment for genes involved in the immune response. These results are consistent with the reported microglial activation in AD^56^, PD^57^and ASD^58^ and also with the crucial role of microglia in the CNS development and immunity^59,60^. The astrocyte-specific module (M5), broadly upregulated in AD, PD, PA, ASD, Scz, and BP (**Fig. 5a**), was enriched for metabolic genes. An increase in astrocytic reactivity has previously been reported in response to oxidative stress induced by amyloid-beta accumulation^61^. Additional analysis demonstrated significant overlap of module M5 with an astrocyte-specific module (CD4: yellow) from Gandal et al.^21^ study which represented coordinated upregulation in ASD, BP and Scz (**Fig. S19c**). These results highlight the significant role of astrocytes in synaptic signalling, neuroprotection, and brain development^62–64^.

Furthermore, we investigated the role of enhancers in the regulation of the network modules. We used an independently derived dataset of brain enhancer RNAs or eRNA (a class of relatively long non-coding RNAs) modules^65^ (sets of eRNAs which are coordinately expressed in the brain). Results demonstrated the enrichment of multiple eRNA modules for the cell-type-specific modules (**Fig. 5c & Supplementary Fig. S20**), demonstrating the role of these enhancer modules in regulating network modules across brain conditions. Neuron-specific modules M8 and M13 showed enrichment for eRNA eModule5, which shows expression specificity for cerebral cortex^65^. In addition, the relationship between protein-coding genes and their adjacent lncRNAs in neuron modules show their coordinated regulation across the conditions (**Fig. 5d**). Oligodendrocyte and microglia modules (M2 and M10) were enriched for eModule7 which is specific for the thalamus-a region affected in a variety of systemic or metabolic diseases, degenerative disease, and psychiatric conditions^66^. The astrocyte-specific module M5 is associated with eModule6, which is specific for medulla oblongata^65^ (a part of the brainstem), a region associated with neurodegeneration and movement disorders^67–69^. These results demonstrate regulation of co-expression modules across the conditions in the brain.

Eventually, as mitochondrial genes have been formerly linked to neuronal phenotypic diversity and brain conditions^70–73^, we performed enrichment analysis of synaptic and nonsynaptic mitochondrial genes for each module using an independent dataset^73^. Multiple modules (M15, M13, M8, M4, and M3) were enriched for mitochondrial genes^73^, of which M15, M8 and M3 showed enrichment for synaptic mitochondria (**Fig. 5e & Supplementary Data 7**). These modules which have previously shown enrichment for neurons (**Fig. S19a**) are mainly downregulated across conditions (**Fig. 5a**). These results highlight the relationship between alteration of mitochondrial transcriptome and synaptic dysfunction in the brain across conditions^74^.

## DISCUSSION

Leveraging the transcriptome profile of post-mortem tissues from several brain regions, for the first time to our knowledge, we highlighted the substantial overlapping molecular patterns across eight brain conditions including NDs and NPs. Dysregulation of overlapping genes such as *SERPINA3, Socs3, APOL4*, and *NPAS4* across brain regions suggests shared perturbation of several mechanisms such as activation of microglia^75^, inflammatory mechanisms, synapse development, and synaptic plasticity^52^ across conditions. Microglia and astrocytes are vital in regulating neuronal activity and brain functioning during development and in the adult brain^76^. These results are consistent with the previous well-established findings that molecular mechanisms in microglia and astrocytes are altered in ND-NPs ^64,77^.

In line with this, co-expression network results revealed mainly downregulation of neuron-specific modules across multiple conditions, reflecting neural dysfunction in both NDs^78^ and NPs^79^. The microglial-related module showed mainly upregulation in NDs (i.e., AD and PD) reflecting activation of microglia during neurodegeneration and brain dysfunction^59,80^. Astrocyte-and oligodendrocyte-specific modules demonstrated broad upregulation across conditions including both NDs and NPs, representing activation of these cell types in neurogenesis, signaling, and cell development^21,62,81^. Also, coordinated downregulation of synaptic mitochondria-related modules across conditions suggests the importance of mitochondria for synaptic connections, neuronal survival, and function^73^ in both NDs^82^ and NPs^83^.

Moreover, our results revealed shared transcriptional changes between neurodegenerative and psychiatric disorders. Specifically, we observed similar transcriptional changes between PD-ASD, AD-ASD, PD-BP, PD-Scz, and AD-Scz. Moreover, within NDs, we found transcriptome alteration overlaps between AD-PD and PA-PSP, while within NPs, we found top transcriptome alteration similarities between Scz-BP and Scz-ASD. The high correlations observed for the Scz-BP and Scz-ASD pairs support previous reports on the molecular similarity of these disorders^21,22^. Correlation of transcriptomic alterations across brain regions demonstrated the limbic lobe captures the majority of transcriptomic similarity between Scz and BP, supporting the role of this region in mood and psychotic disorders^84^. Although the only region studied for PD was the frontal lobe, similar transcriptional changes were observed between PD-ASD, particularly for the genes involved in pathways such as neuroinflammation that has been recently linked to autism^85,86^. High frequency of parkinsonism has previously been reported in autistic cases^87^, in which inflammatory mechanisms are seemingly involved in the pathobiology of the disease^88,89^. The correlation of transcriptional alterations between AD and ASD supports the evidence of neurodegeneration in autism^90^. This similarity between AD and ASD was mainly observed for the temporal and then frontal lobes. The overlapping expressional changes of genes such as *CHI3L1* between Scz and PD suggest perturbation of dopaminergic-glutamatergic balance in the brain, as described previously^91,92^. The Scz-AD similarity, which was mainly captured by the temporal lobe, is also consistent with the evidence of shared mechanisms between neurodegeneration and Scz^93,94^. Primary damaged regions in PSP and PA are reported to be brain stem (specifically substantia nigra)^95^ and hippocampus^96^, respectively. However, the significant correlation of transcriptome alterations between PSP and PA, mainly observed in the temporal lobe, suggests similar molecular changes of this brain area in the conditions. Although PSP is sometimes misdiagnosed as PD due to similar clinical symptoms^97^, we did not observe transcriptome similarity here. This could be due to the lack of primary damaged regions for either condition and/or different pathobiology of the conditions^98^.

In addition, our findings provide new insights about the shared cognitive impairment in ASD and Scz and their transcriptome similarity with NDs^99,100^. Also, the results did not show significant transcriptional correlation between AD and PA, suggesting their divergent molecular pathobiology^101^. Transcriptional relationships were not observed between MDD and other NPs such as Scz and ASD as expected, which could be due to its heterogeneous nature^21,22,102^. Of note, shared transcriptional changes observed here do not necessarily indicate similar pathobiology of the conditions, particularly in conditions such as PD, PSP, and PA that lacked primary damaged regions which demands further investigation.

Comparisons of transcriptional changes across brain regions demonstrated similar molecular changes in the temporal and frontal lobe across NDs and NPs, implicating their possible impairment in the pathogenesis of a variety of brain diseases^103–109^. Similar molecular patterns of the cerebellum were observed across AD, PSP, and PA, supporting its emerging role in the pathobiology of NDs^110,111^. Despite the lack of samples for some of the conditions mentioned before, the basal ganglia and limbic lobe showed transcriptional similarities across Scz, ASD, BP and MDD, implying their involvement in several mood and psychiatric disorders. These findings suggest, on one hand, the impairment of multiple brain regions (rather than one primary region) in one condition and, on the other hand, the involvement of one region in multiple conditions. Of note, in the analysis of the overlapping transcriptomic alterations, we did not expect to capture modifications directly linked to the underlying etiological mechanisms of the different conditions studied here. However, this was addressed in condition-specific gene expression analyses and will be more investigated in future analyses with the same database, along with the exploration of conditions sharing common mechanisms (i.e., cerebral proteinopathies) or between the different stages of the same disease (i.e., preclinical and clinical stages of AD)^112,113^. In line with this, we anticipate that future research will also benefit from the integration of transcriptomics with other omics modalities, such as genomics, proteomics, metabolomics, and epigenomics. This promises to provide deeper insights into the causative pathways through which genes and environment interact during life and influence the human brain^114^. Additional research could also benefit from further identification of sex-specific gene networks and transcription profiles to unravel the molecular mechanisms of brain diseases^115–117^.

In addition, since tissue samples from all brain regions were not available for all conditions, in some of them we might have missed the transcriptomic profile of the brain area in which the primary pathology is expected to be expressed (i.e., basal ganglia in PD). Also, even though batch-effect corrections have thoroughly been applied here, the inevitable effect of merging multiple datasets on the results could still be a limitation. However, despite the limitations of the current analyses, molecular signatures here described across ND-NPs can provide target leads for the development of therapeutic interventions targeted to common pathological mechanisms which may overcome indications solely based on clinical manifestations, thus paving the way for the rational design of personalized and mechanistically-based therapies^118^.

## METHODS

### Samples and raw data

RNA-Seq raw data were obtained from 4,711 post-mortem brain samples from subjects with AD, PD, PSP, PA, Scz, MDD, ASD, BP, and matched controls through previously-published studies^8,21,24–41^ and consortia including CommonMind Consortium and PsychENCODE Consortium from Sage Synapse (https://www.synapse.org/) and the NCBI Gene Expression Omnibus (GEO) (https://www.ncbi.nlm.nih.gov/geo/) (see **Table S1**). The samples from individual datasets were processed separately and analyzed according to our RNA-Seq pipeline as described below.

### Data processing

The data obtained from individual datasets were processed separately. We used RNA-Seq *fastq* files as the initial source of data processing. The samples that were retrieved as *SRA* and *BAM* files were converted to *fastq* file formats using SRA-toolkit^119^ and SAM-tools^120^, respectively. For further sample processing, the Grape pipeline^121^ was used for RNA-Seq analysis, with Nextflow^122^ as the execution backend, the STAR aligner v.2.6.0a tool^123^ for mapping reads to the human genome build hg19 with GENCODE v.28 annotations, and the RSEM tool^124^ for isoform quantification (using default options). Next, post-alignment quality control (QC) was performed using STAR aligner statistics, Qualimap v.2.2.1 tools^125^, and Picard tools v1.8 (http://broadinstitute.github.io/picard/) to check for the total number of reads, total number of mapped reads, GC percentage, exonic rate, intronic rate, intergenic rate, duplication rate, and insertion/deletion rate. Sequencing statistics was used to check for quality control and sample outliers. The data were then normalized for library size using the *voom-limma* R package^126^. To filter out lowly-expressed genes, only genes with at least log_2_(CPM) of 0.5 in 70 % of the samples were kept for further analyses. The *sva* R package v.3.32.1^127^ was used to correct for any batch effect of sequencing library preparations. To remove sample outliers, standardized network connectivity Z-scores were measured and a cutoff of Z < −2 was set as the threshold^128,129^. Normalized datasets were kept for downstream analyses.

### Samples clustering and tSNE analysis

t-SNE^43^ was used to produce a latent representation of the samples based on their gene expression profiles across datasets using all normalized datasets which were pooled together in an expression matrix. Before this, principal component analysis (PCA)^130^ was used to reduce the number of dimensions and obtain a small number of principal components as input to the tSNE analysis. The first two t-SNE coordinates were used for visualization.

### Differential Gene Expression (DGE) analysis and transcriptome comparisons

Differential expression analysis for each dataset was performed using *limma* with empirical Bayes moderated t-statistics, with the following model (expression ∼ diagnosis + age + sex + RIN + ethnicity + PMI + pH). For condition- and brain region-specific DGE analyses, normalized expression data from relevant studies were combined and DGE was calculated using linear mixed-effects models by the *nlme* R package v.3.1-140^131^, with fixed effects of diagnosis, age, sex, and study (and brain region when calculating condition-specific DE). A random effect for subjects was used to fix for overlapping subjects between the studies (expression ∼ diagnosis + age + sex+ study + 1 | subject). The calculated log fold-change (logFC) values were used for downstream analyses. Significantly differentially expressed genes (DEGs) were filtered by using a p-value of < 0.05 and |log_2_FC| > 0.5 as a significant threshold.

### Reproducibility of differential expression results

In order to check reproducibility of the results from DGE, the list of DEGs obtained from each dataset was compared to those obtained from condition-specific analysis by calculating Jaccard index^132^ and Simpson’s index^133^ using *GeneOverlap* R package^134^. Fisher’s exact test was used to calculate p-values of each comparison.

### Building classifier models

To identify prediction power of transcriptional alterations from frontal and temporal lobe between cases and controls, normalized expressions of DEGs for each region were used to build classifier models using random forests^135^. Accuracy, sensitivity and specificity of the final models were used for comparing the results. A 5-fold cross-validation was used to avoid overfitting as much as possible.

### Comparisons of transcriptional alterations

To analyze cross-condition transcriptome profile comparisons, we only kept the 15,819 genes that were common across all diseases. Pairwise gene expression comparisons were performed using Spearman’s correlation over logFC values of the genes. In addition, a correlation network and tree dendrogram was built using the correlations statistics to obtain a relative relationship of the conditions based on transcriptional alterations. To check reproducibility of results, logFC values of the genes common across condition-specific analyses and individual datasets were compared using Spearman’s correlation test. Moreover, brain region-specific comparisons across conditions were performed using logFC values of the genes shared between the present conditions for each region.

### Rank-Rank Hypergeometric Overlap (RRHO) analysis

In order to highlight the degree of overlap in gene signatures across conditions, as well as comparing condition pairs for shared brain regions, we performed an unbiased rank-rank hypergeometric overlap (RRHO) analysis using the *RRHO* R package v.1.24.0^136^. A one-sided version of the test only looking for over-enrichment was used. RRHO difference maps were produced by calculating for each pixel the normal approximation of difference in the log odds ratio and standard error of overlap with expression data in the intersection list. P-values were calculated and FDR-corrected for multiple comparisons across pixels.

### Gene co-expression network analysis

We performed robust Weighted Gene Co-Expression Network Analysis (rWGCNA) using the *WGCNA* R package v.1.68^53^ to identify co-expressed gene modules using expression data that were first normalized for different covariates. The expression datasets from independent condition-specific DGE analyses were combined using the 15,819 genes common between all datasets. Batch effect correction for the studies was performed using the *ComBat* function from the *sva* R package.

Co-expression analysis was then performed using signed networks. Co-expression networks were built using the *blockwisemodules* function. The network dendrogram was created using the “average” linkage hierarchical clustering of the topological overlap dissimilarity matrix to identify modules of highly co-regulated genes. To obtain an approximately scale-free weighted co-expression network, a power function with a soft-threshold of 20 was applied to the merged expression dataset. Modules were defined as branches of the dendrogram using the hybrid dynamic tree-cutting method, followed by a dynamic cut-tree algorithm to separate clustering dendrogram branches into gene modules. Modules were then summarized by their first principal component (ME, module eigengene) and those with high eigengene correlations were again merged.

Because the topological overlap between two genes reflects both their direct and indirect interactions through all other genes in the network, this approach helps to build more cohesive and more biologically meaningful modules. To ensure the robustness of the module, random resampling was performed from the initial set of samples 100 times followed by consensus network analysis. The final module was achieved using network parameters including biweight midcorrelation (bicor), a minimum module size of 50, deepsplit of 4, merge threshold of 0.1, and negative pamStage. Each module was assigned by a unique (and arbitrary) color identifier. Genes with the highest intramodular connectivity (those with more connections at the core of the network) were considered as hub genes. Significance values were FDR-corrected to account for multiple comparisons. Top hub genes with the most connections were prioritized based on their module membership (kME), defined as a correlation to the module eigengene. Module-condition relationships were measured using a linear mixed effects model, with a random effect for subjects to fix for overlapping subjects.

### Cell-type-specific enrichment analysis

To analyze cell-type-specific gene expression within each module, we retrieved the single-cell data for human brain cell types from the PanglaoDB database^54^. The genes within each module were then compared to the marker genes for each brain cell type using the *GeneOverlap* R package v.1.20.0^134^. Fisher’s exact test with an FDR-correction for p-values was used to analyze the gene overlap comparisons. To check the consistency of the results, another cell-type-specific expression dataset composed of five main brain cell types including neurons, astrocytes, oligodendrocytes, microglia, and endothelial cells was obtained from another single-cell RNAseq study^55^. Gene symbols were mapped to Ensembl gene identifiers using the *biomaRt* R package. Specificity for the five brain cell types was calculated with the *specificity*.*index* function from *pSI* R package v.1.1 ^137^. Fisher’s exact test with FDR correction for p-values was applied to check for the significant cell-type specificity (FDR-corrected p-value <0.05 was considered statistically significant).

To see transcriptional relationships of cell-type-specific modules across conditions, effect sizes (beta values) obtained from linear mixed effect models for genes within each module were compared using Spearman’s correlation, followed by FDR-correction for multiple comparison tests.

### Gene ontology enrichment analysis

Gene Ontology (GO) pathway enrichment for each condition and gene modules was performed using the *gprofiler2* R packages. Only pathways that comprise 10 to 2000 genes were filtered for analysis. The top pathways with an FDR-corrected p-value < 0.05 were considered significantly related.

### Brain enhancer RNAs co-expression analysis

To understand the relationship between regulatory factors in the brain and co-expression modules, an expression dataset for brain enhancer-RNAs (eRNA)-a non-coding RNA transcribed from active enhancers^65^-was obtained from an independent study of human brain region-specific eRNAs co-expression analysis^65^. To explore the co-expression of gene modules from our dataset and each brain eRNA module, overlap of genes within each module and genes from eRNA modules was tested by Fisher’s exact test. A p-value of < 0.05 followed by FDR correction was used to filter the significant enrichments.

### Protein coding-lncRNA co-expression analysis

Within each co-expression module, genomic coordinates for the genes were obtained using the *BiomaRt* R package. Next, the genes were filtered to protein-coding genes and lncRNA pairs with a distance of < 10Mb, as the *cis*-regulatory cutoff distance. The expression fold change (beta) values of the genes across the conditions were illustrated in a genomic circos plot using the *circlize*^*138*^ R package.

### Mitochondrial transcriptome enrichment analysis

To see whether co-expression modules are enriched for mitochondrial transcriptome, an independent dataset containing synaptic and nonsynaptic mitochondria modules was obtained^73,139^. Enrichment analysis for each module was performed by Fisher’s exact test followed by FDR correction for p-values.

### Software and code availability

The R programming language version 3.5.0 (https://www.r-project.org/) was used for statistical analyses and data visualization. The functions and libraries used in this study are available as R packages: *WGCNA, nlme, RRHO, GeneOverlap, pSI, ggplot2, Rtsne, gprofiler2, caret, limma, pheatmap, ComplexHeatmap, circlize* at CRAN (http://cran.r-project.org/) and/or Bioconductor (https://bioconductor.org/).

## Data availability

The raw data incorporated in this work were gathered from various resources. RNA-Seq raw data, metadata, and source files are available on the NCBI GEO database and Sage Synapse as described in **Supplementary Table S1**.

## Acknowledgements

RNA-seq data (retrieved from Sage Synapse with accession number syn2759792, with access governed by NIMH Repository and Genomics Resource) were generated as part of the PsychENCODE Consortium supported by U01MH103339, U01MH103365, U01MH103392, U01MH103340, U01MH103346, R01MH105472, R01MH094714, R01MH105898, R21MH102791, R21MH105881, R21MH103877, and P50MH106934 awarded to Schahram Akbarian (Icahn School of Medicine at Mount Sinai), Gregory Crawford (Duke), Stella Dracheva (Icahn School of Medicine at Mount Sinai), Peggy Farnham (USC), Mark Gerstein (Yale), Daniel Geschwind (UCLA), Thomas M. Hyde (LIBD), Andrew Jaffe (LIBD), James A. Knowles (USC), Chunyu Liu (UIC), Dalila Pinto (Icahn School of Medicine at Mount Sinai), Nenad Sestan (Yale), Pamela Sklar (Icahn School of Medicine at Mount Sinai), Matthew State (UCSF), Patrick Sullivan (UNC), Flora Vaccarino (Yale), Sherman Weissman (Yale), Kevin White (UChicago) and Peter Zandi (JHU).

RNA-seq data (retrieved from Sage Synapse with accession number syn4921369 governed by NIMH Repository and Genomics Resource) were generated as part of the CommonMind Consortium supported by funding from Takeda Pharmaceuticals Company Limited, F. Hoffmann-La Roche Ltd and NIH grants R01MH085542, R01MH093725, P50MH066392, P50MH080405, R01MH097276, RO1-MH-075916, P50M096891, P50MH084053S1, R37MH057881, AG02219, AG05138, MH06692, R01MH110921, R01MH109677, R01MH109897, U01MH103392, and contract HHSN271201300031C through IRP NIMH. Brain tissue for the study was obtained from the following brain bank collections: the Mount Sinai NIH Brain and Tissue Repository, the University of Pennsylvania Alzheimer’s Disease Core Center, the University of Pittsburgh NeuroBioBank and Brain and Tissue Repositories, and the NIMH Human Brain Collection Core. CMC Leadership: Panos Roussos, Joseph Buxbaum, Andrew Chess, Schahram Akbarian, Vahram Haroutunian (Icahn School of Medicine at Mount Sinai), Bernie Devlin, David Lewis (University of Pittsburgh), Raquel Gur, Chang-Gyu Hahn (University of Pennsylvania), Enrico Domenici (University of Trento), Mette A. Peters, Solveig Sieberts (Sage Bionetworks), Thomas Lehner, Stefano Marenco, Barbara K. Lipska (NIMH).

## Author contributions

I.S., V.D., R.G., and N.V.-T. conceived the study. I.S. performed the analyses. E.P., M.M.-A., G.S., A.N., R.G., and N.V.-T. provided statistical and analysis advice. J.D.G., V.W., M.M.-A., G.S., A.N., R.G., and N.V.-T. contributed to the interpretation of the results. I.S., J.D.G., R.G. and N.V.-T. wrote the manuscript. R.G. and N.V.-T. supervised the project. All authors read and approved the final manuscript.

