## Supplementary material for "Brain transcriptomic profiling reveals common alterations across neurodegenerative and psychiatric disorders": Supplementary document.pdf (from another version of this preprint: version 1, 2021-08-16)

**Table S1.** RNA-Seq expression datasets obtained to use in this study.

| Diagnosis | Study/Project | Accession ID | # Samples | | Brain region | Ref |
|---|---|---|---|---|---|---|
| | | | Controls | Cases | | |
| **AD** | Wang et al | syn3159438 | 280 | 742 | Anterior prefrontal cortex, perirhinal cortex, superior temporal gyrus, pars opercularis | [1] |
| | Allen et al | syn5550404 | 155 | 164 | Temporal cortex, cerebellum | [2] |
| **PD** | Dumitriu et al | PRJNA283498 | 44 | 29 | BA9 | [3] |
| **PA** | Allen et al | syn5550404 | Mentioned (155) | 58 | Temporal cortex, cerebellum | [2] |
| **PSP** | Allen et al | syn5550404 | Mentioned (155) | 168 | Temporal cortex, cerebellum | [2] |
| **Scz** | Jaffe et al | syn12299750 | 320 | 175 | Dorsolateral prefrontal cortex (DLPFC) | [4] |
| | Xiao et al | PRJNA235930 | 12 | 10 | BA9, BA24 | [5] |
| | Chang et al | PRJNA379666 | 24 | 22 | Amygdala | [6] |
| | Corley et al | PRJNA343829 | 19 | 19 | DLPFC | [7] |
| | Wu et al | PRJEB2939 | 9 | 9 | Superior temporal gyrus | [8] |
| | CommonMind consortium | syn18097439 | 294 | 47 | DLPFC | [9] |
| | CommonMind consortium-HBCC | syn18097439 | 167 | 87 | DLPFC | [9] |
| | BrainGVEX | syn4590909 | 257 | 95 | Frontal cortex | [10] |
| | Ramaker et al | PRJNA319583 | 70 | 71 | anterior cingulate cortex, DLPFC, | [11] |

| | | | | | | |
|---|---|---|---|---|---|---|
| | | | | | nucleus accumbens | |
| **ASD** | Wright et al | PRJNA398545 | 39 | 13 | DLPFC | [12] |
| | Li J et al | PRJNA263196 | 6 | 6 | Corpus callosum | [13] |
| | Liu et al | PRJNA254971 | 38 | 34 | Superior frontal gyrus | [14] |
| | Yale-ASD | syn4566141 | 30 | 15 | DLPFC | [10] |
| | UCLA-ASD | syn4587609 | 126 | 119 | BA4/6,<br>BA38,<br>BA7,<br>BA17 | [10] |
| **MDD** | Ramaker et al | PRJNA319583 | Mentioned (70) | 69 | anterior cingulate cortex,<br>DLPFC,<br>nucleus accumbens | [11] |
| | Pantazatos et al | PRJNA394722 | 29 | 30 | DLPFC (BA9) | [15] |
| | Labonte et al | PRJNA398031 | 122 | 141 | Orbitofrontal cortex (BA11),<br>DLPFC (BA8/9),<br>cingulate gyrus 25 (BA25; vmPFC),<br>anterior insula,<br>nucleus accumbens,<br>ventral subiculum | [16] |
| **BP** | Xiao et al | PRJNA235930 | Mentioned (12) | 14 | BA9,<br>BA24 | [5] |
| | CommonMind consortium | syn18097439 | Mentioned (294) | 257 | DLPFC | [9] |
| | CommonMind consortium-HBCC | syn18097439 | Mentioned (167) | 58 | DLPFC | [9] |
| | BrainGVEX | syn4590909 | Mentioned (257) | 73 | Frontal cortex | [10] |
| | Ramaker et al | PRJNA319583 | Mentioned (70) | 71 | anterior cingulate cortex,<br>DLPFC,<br>nucleus accumbens | [11] |
| | Pacifico et al | PRJNA318642 | 18 | 18 | Dorsal striatum | [17] |

| | | | | | | |
|---|---|---|---|---|---|---|
| | McMullen et al | PRJNA321439 | 8 | 8 | Putamen, Caudate nucleus | [18] |
| | Akula et al | PRJNA231202 | 11 | 11 | DLPFC | [19] |

**Table S2.** Number of studies per region/condition

| | cerebellum | Temporal | Frontal | Limbic | Occipital | Insular | Basal ganglia |
|---|---|---|---|---|---|---|---|
| AD | 1 | 2 | 1 | | | | |
| PD | | | 1 | | | | |
| PA | 1 | 1 | | | | | |
| PSP | 1 | 1 | | | | | |
| Scz | | 2 | 8 | 2 | | | 1 |
| ASD | 2 | 1 | 4 | 1 | 1 | | |
| MDD | | | 3 | 2 | | 1 | 2 |
| BP | | | 5 | 2 | | | 3 |

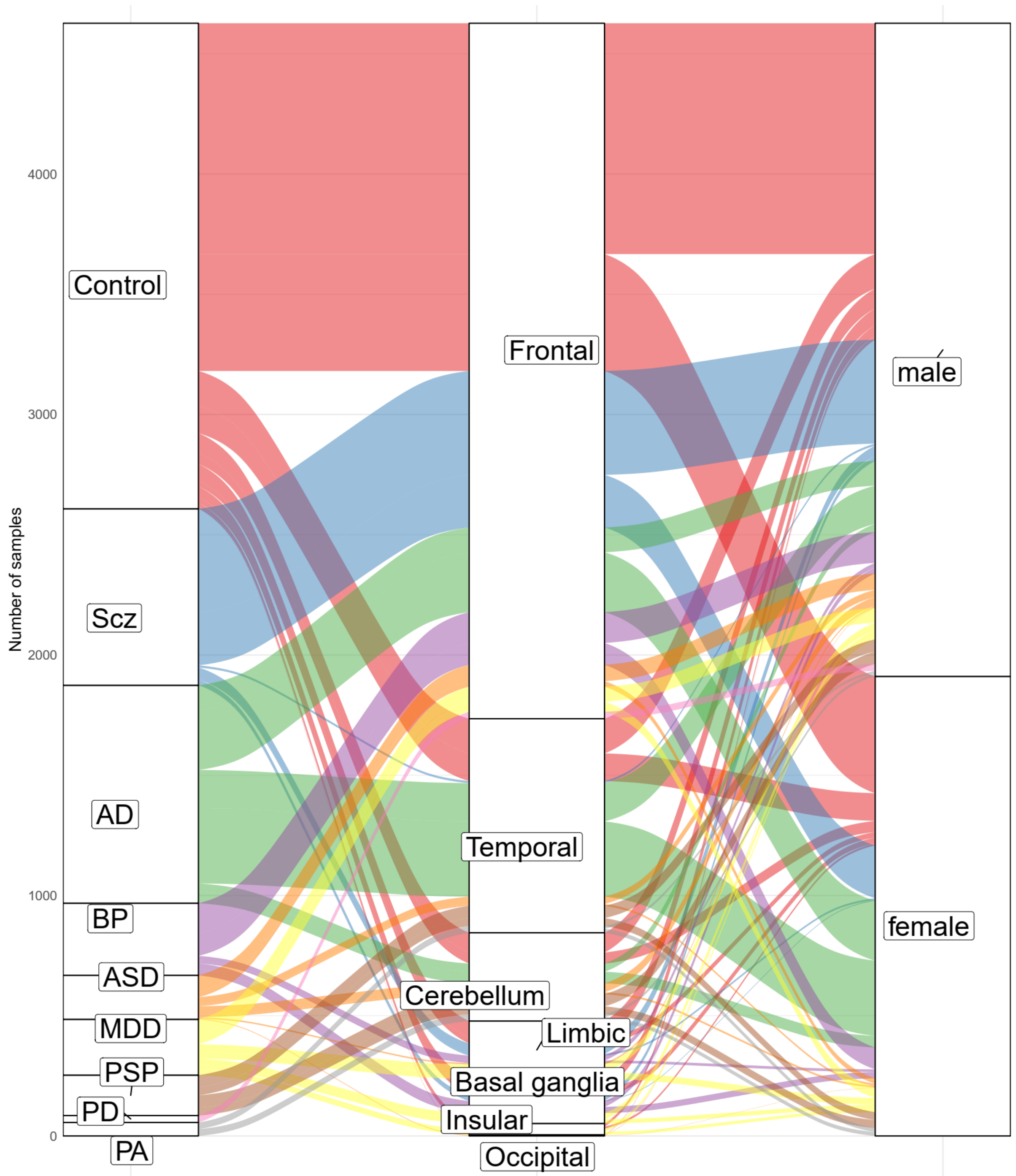


**Figure S1**. The overview of samples used in this study. A flowchart of the samples. From left to right, the panels represent diagnosis, brain lobes, and sex, with colors showing diagnosis.

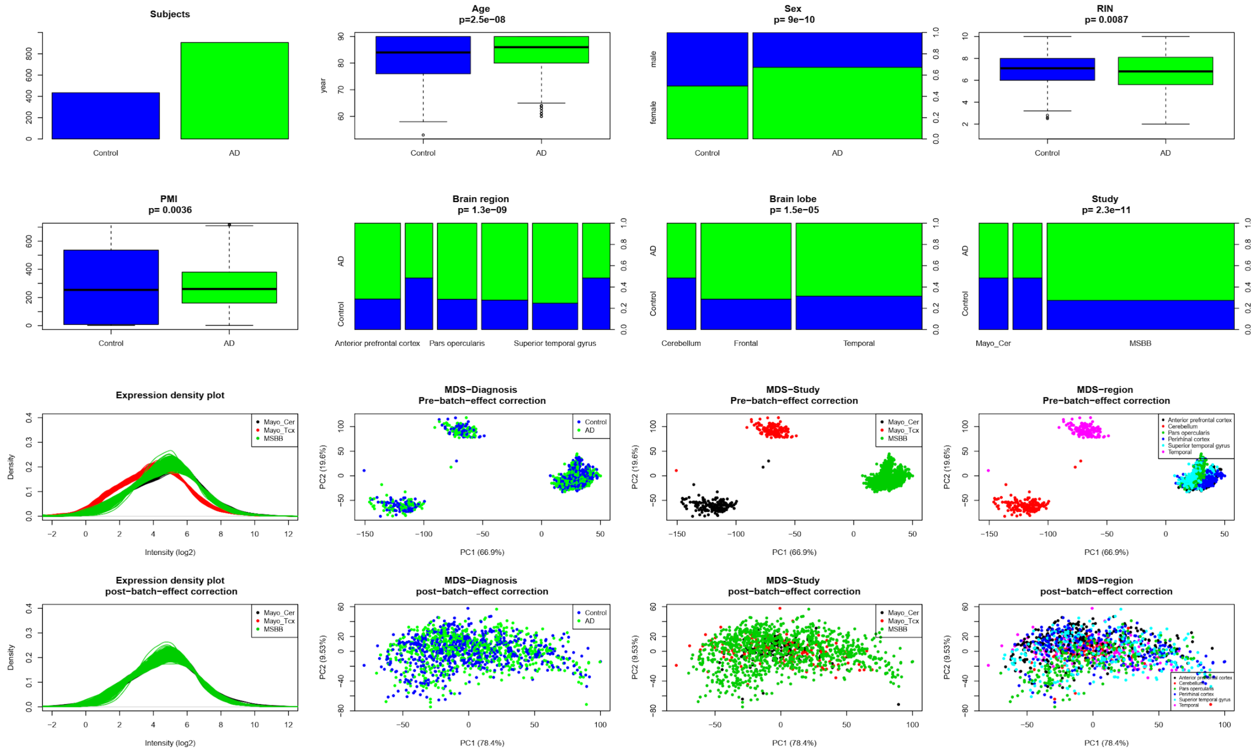


**Figure S2.** Quality control (QC) plots are shown for AD datasets. Normalized data from the cohorts analyzed in this study consisted of the anterior prefrontal cortex, perirhinal cortex, superior temporal gyrus, pars opercularis, temporal cortex, and cerebellum brain samples from subjects with AD (n = 906) and controls (n = 479). Sample outliers were detected by standardized network connectivity z-scores and removed. Batch effects were corrected for the studies. Multidimensional scaling (MDS) plots show sample clustering by the first two expression principal components. Groups were balanced by available covariates and potential confounding factors.

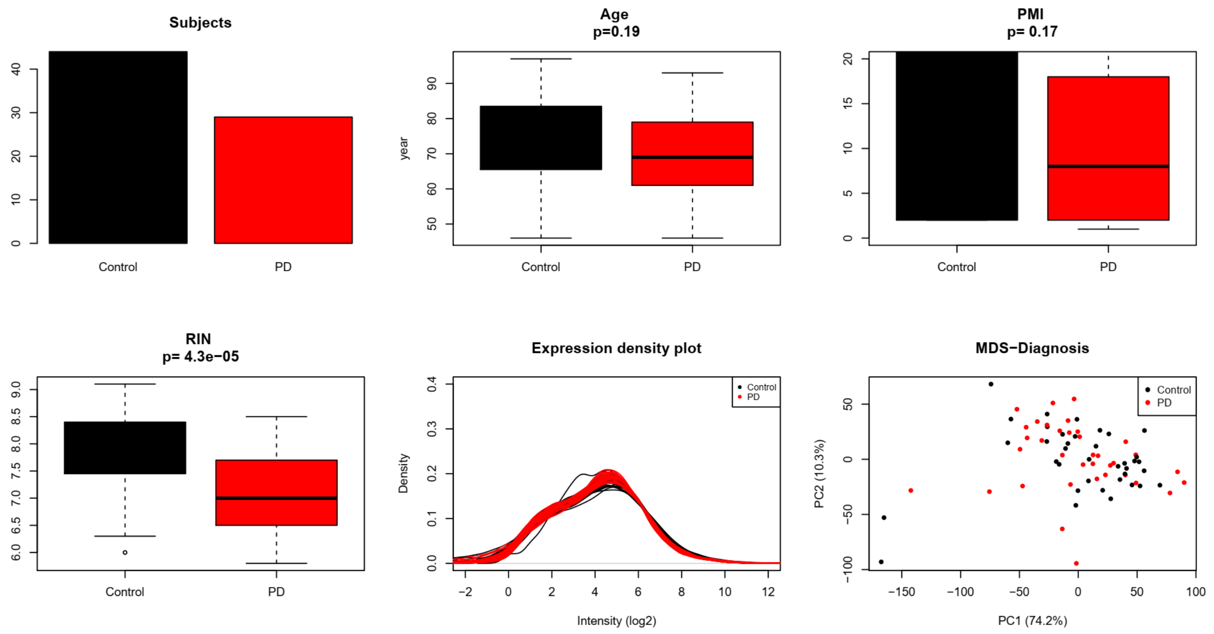


**Figure S3.** QC plots for the PD dataset. This dataset consisted of dorsolateral prefrontal cortex (BA9; frontal lobe) brain samples from subjects with PD (n = 29) and controls (n = 44). Sample outliers were detected by standardized network connectivity z-scores and removed. Multidimensional scaling (MDS) plots show sample clustering by the first two expression principal components. Groups were balanced by available covariates and potential confounding factors.

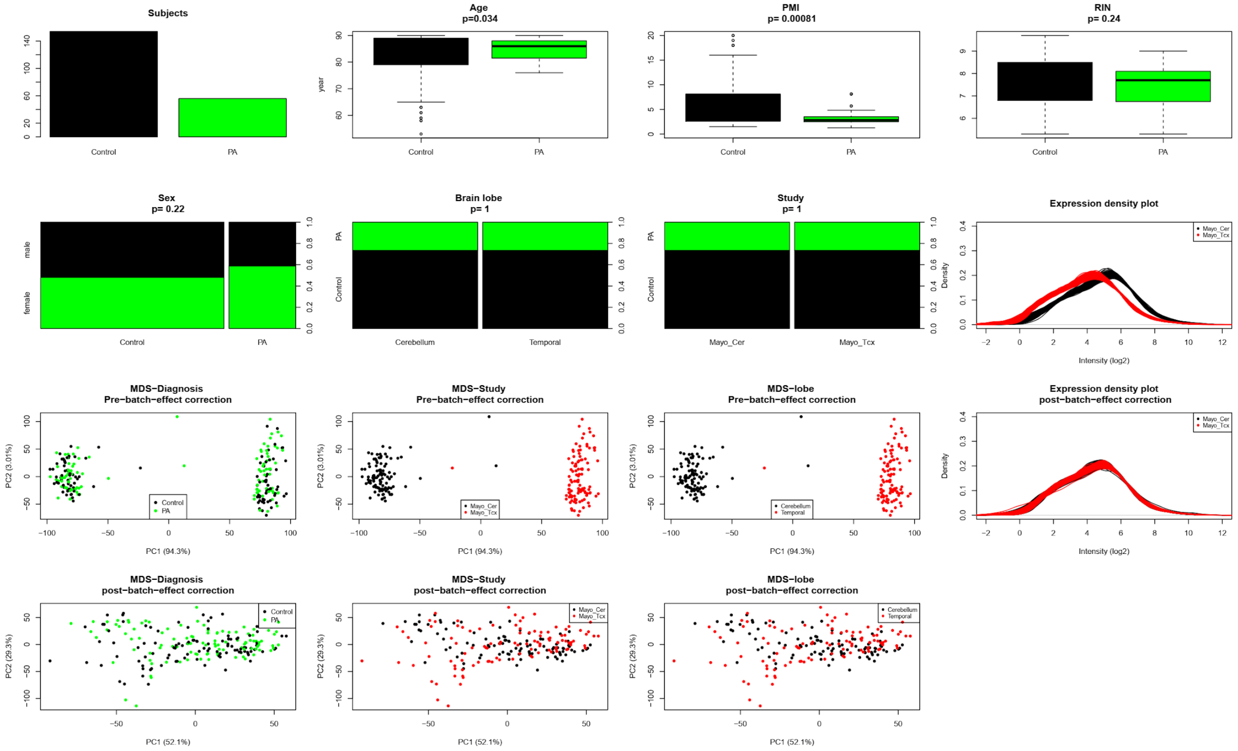


**Figure S4.** QC plots are shown for PA datasets. Normalized data from one group by two cohorts analyzed here consisted of the temporal cortex and cerebellum brain samples from subjects with PA (n =58 ) and controls (n =155 ). Sample outliers were detected by standardized network connectivity z-scores and removed. Batch effects were corrected for the studies. Multidimensional scaling (MDS) plots show sample clustering by the first two expression principal components. Groups were balanced by available covariates and potential confounding factors.

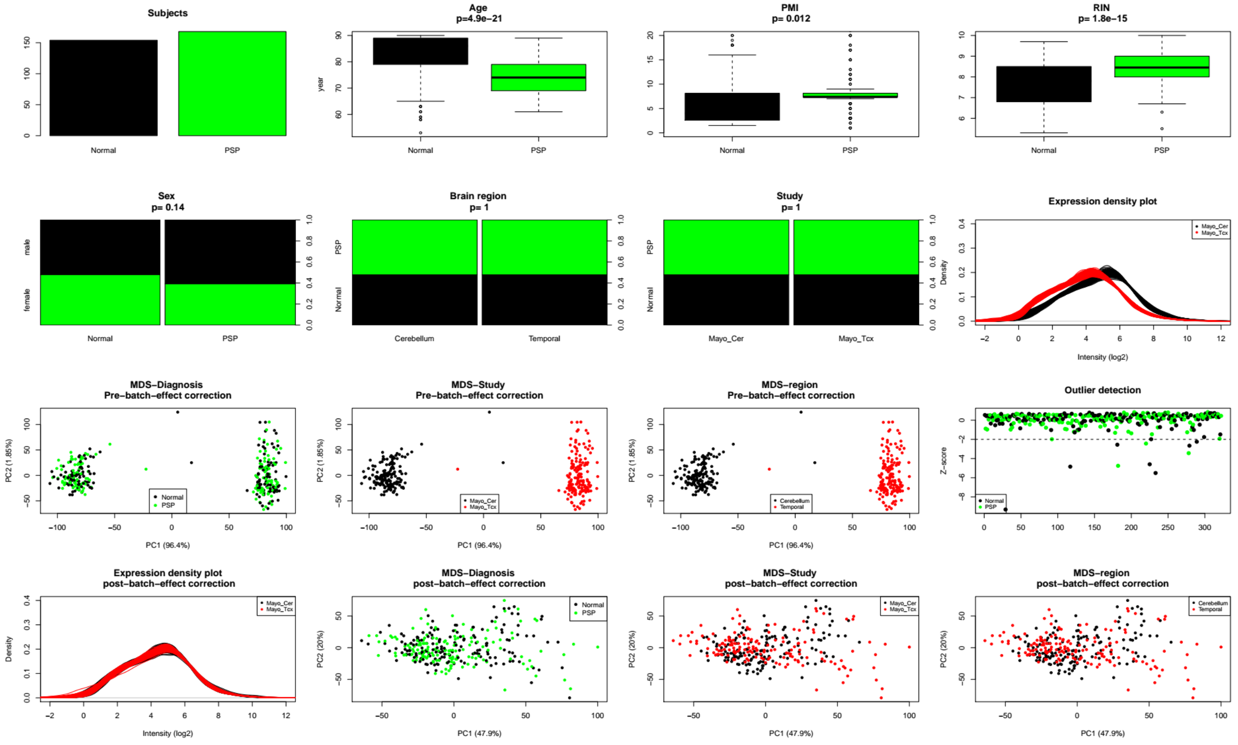


**Figure S5.** QC plots are shown for PSP datasets. Normalized data from one group by two cohorts analyzed here consisted of the temporal cortex and cerebellum brain samples from subjects with PSP (n = 168) and controls (n =155 ). Sample outliers were detected by standardized network connectivity z-scores and removed. Batch effects were corrected for the studies. Multidimensional scaling (MDS) plots show sample clustering by the first two expression principal components. Groups were balanced by available covariates and potential confounding factors.

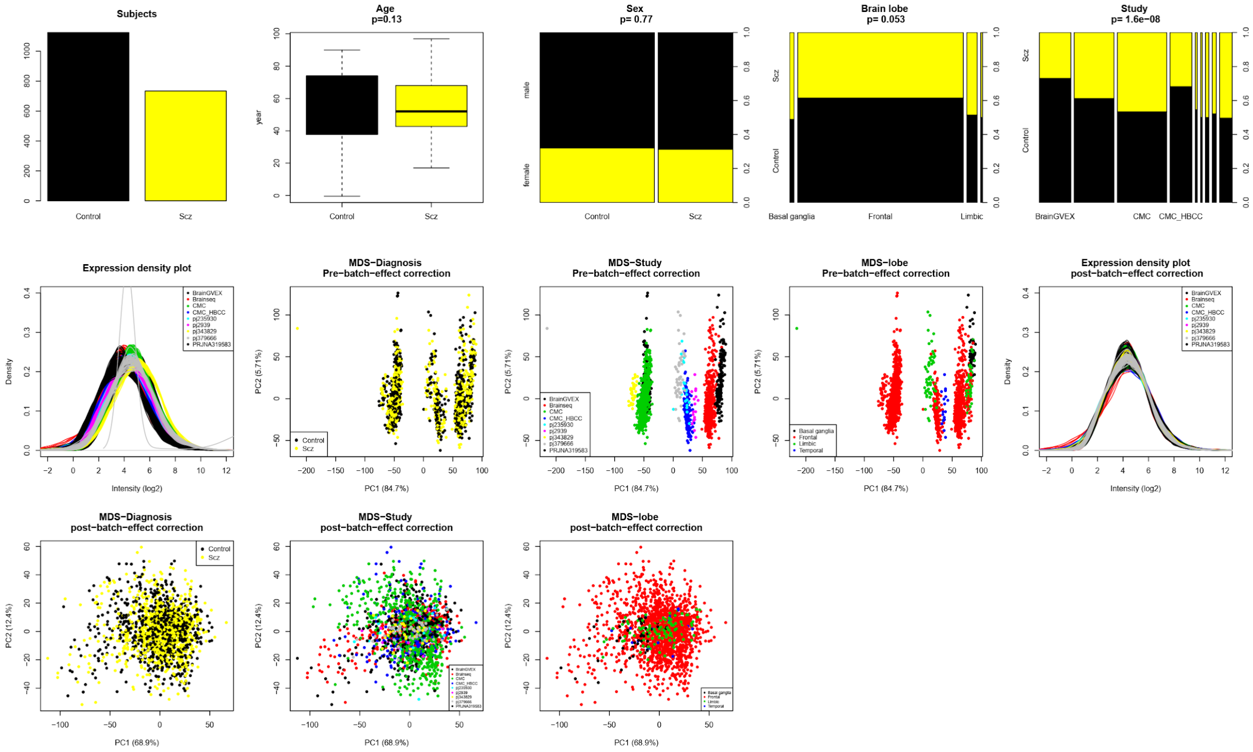


**Figure S6.** QC plots are shown for Scz datasets. Normalized data from nine studies analyzed here consisted of granular frontal area 9, DLPFC, anterior cingulate cortex, ventral anterior cingulate 25, amygdala, superior temporal gyrus, and nucleus accumbens brain samples from subjects with Scz (n = 535) and controls (n = 1172). Sample outliers were detected by standardized network connectivity z-scores and removed. Batch effects were corrected for the studies. Multidimensional scaling (MDS) plots show sample clustering by the first two expression principal components. Groups were balanced by available covariates and potential confounding factors.

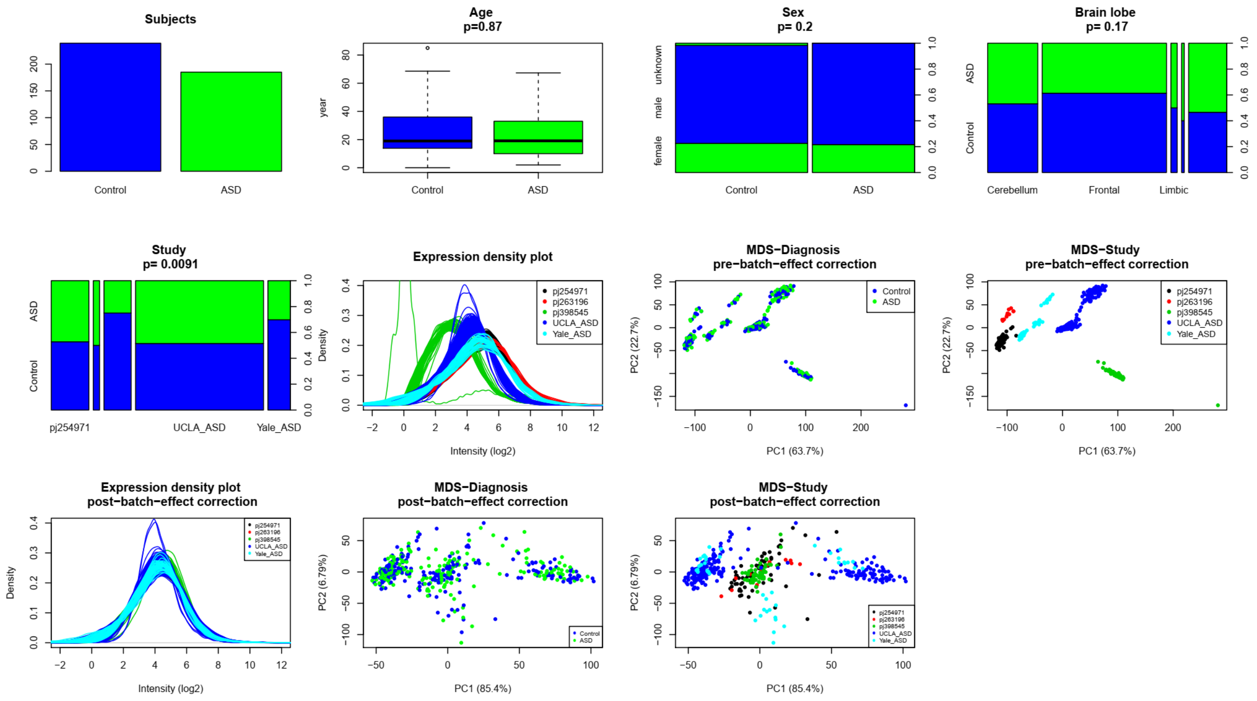


**Figure S7.** QC plots are presented for ASD datasets. Normalized data from five studies analyzed here consisted of the corpus callosum, inferior temporal cortex, auditory cortex, V1C (primary visual cortex), superior frontal gyrus, and DLPFC brain samples from subjects with ASD (n = 187) and controls (n = 239). Sample outliers were detected by standardized network connectivity z-scores and removed. Batch effects were corrected for the studies. Multidimensional scaling (MDS) plots show sample clustering by the first two expression principal components. Groups were balanced by available covariates and potential confounding factors.

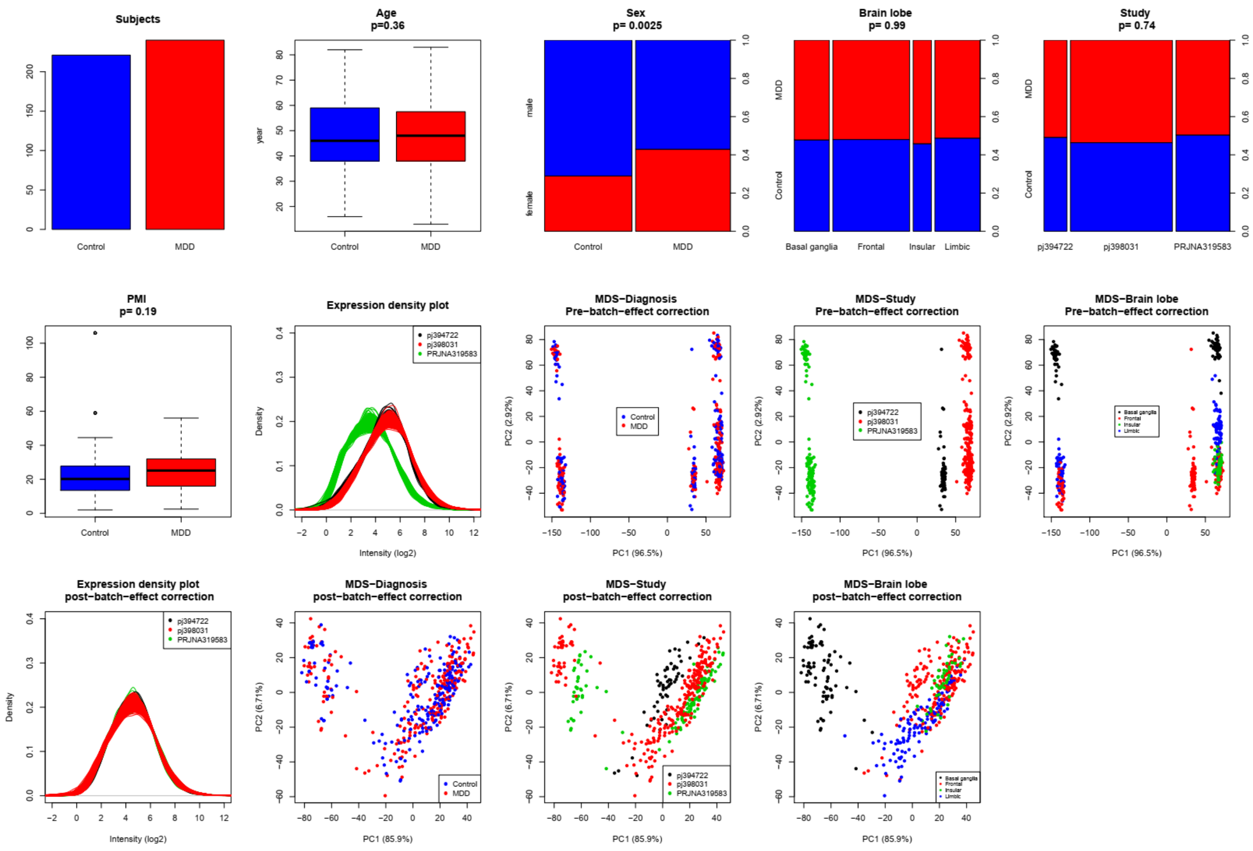


**Figure S8.** QC plots are shown for MDD datasets. Normalized data from three studies analyzed here consisted of the anterior insula (aINS), nucleus accumbens (Nac), DLPFC, orbitofrontal cortex (OFC), ventral subiculum (vSub), cingulate gyrus 25 (Cg25), anterior cingulate cortex, brain samples from subjects with MDD (n = 240) and controls (n = 221). Sample outliers were detected by standardized network connectivity z-scores and removed. Batch effects were corrected for the studies. Multidimensional scaling (MDS) plots show sample clustering by the first two expression principal components. Groups were balanced by available covariates and potential confounding factors.

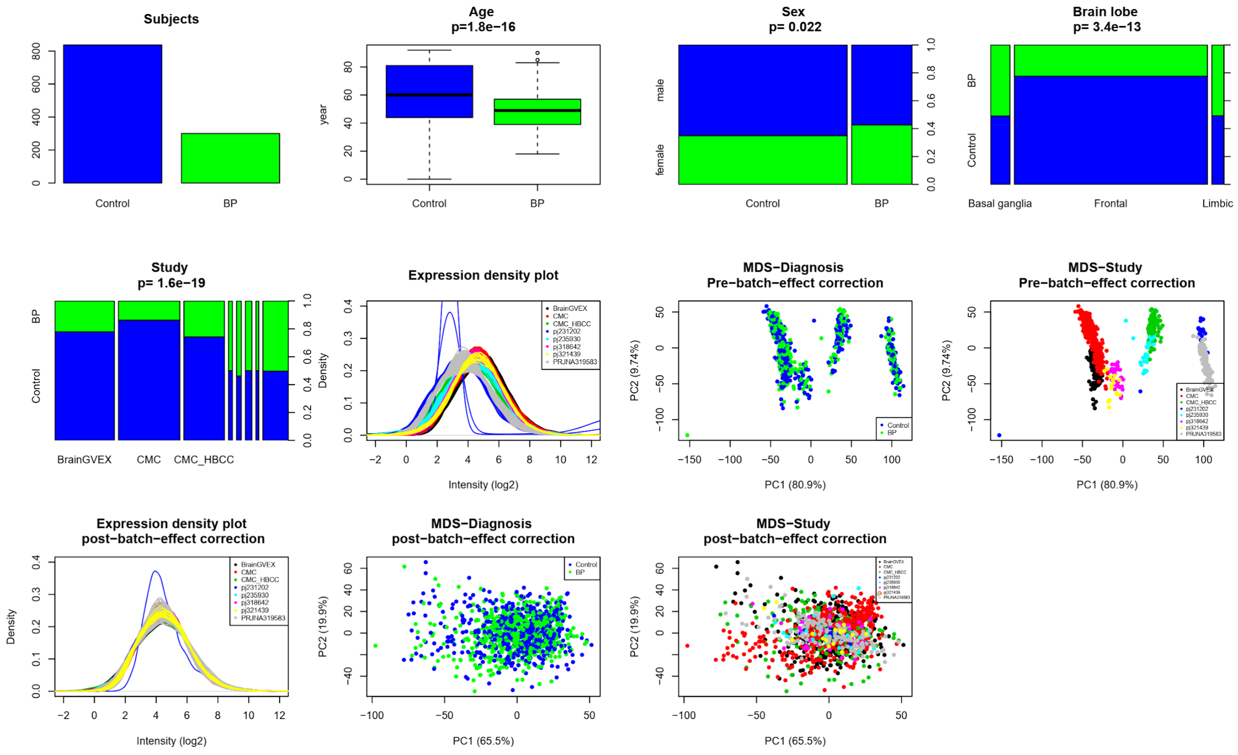


**Figure S9.** A set of QC plots is shown for BP datasets. Normalized data from eight studies analyzed here consisted of the corpus callosum, inferior temporal cortex, auditory cortex, V1C (primary visual cortex), superior frontal gyrus, and DLPFC brain samples from subjects with BP (n = 511) and controls (n = 837). Sample outliers were detected by standardized network connectivity z-scores and removed. Batch effects were corrected for the studies. Multidimensional scaling (MDS) plots show sample clustering by the first two expression principal components. Groups were balanced by available covariates and potential confounding factors.

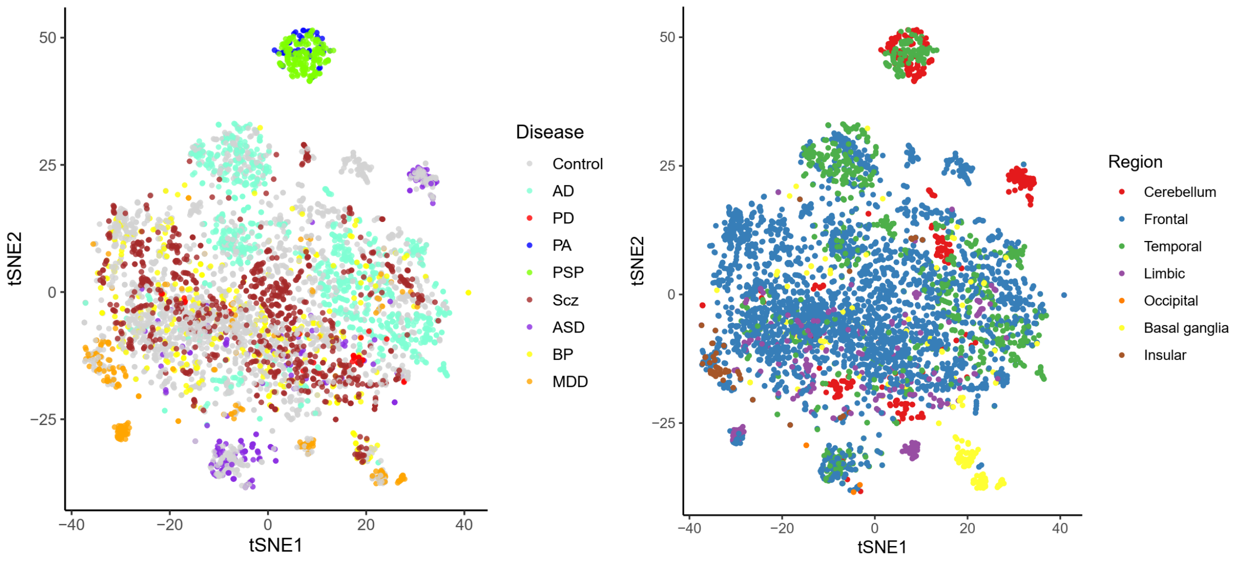


**Figure S10.** tSNE visualization of pooled samples, colored by condition (**left**) and by brain region (**bottom**).

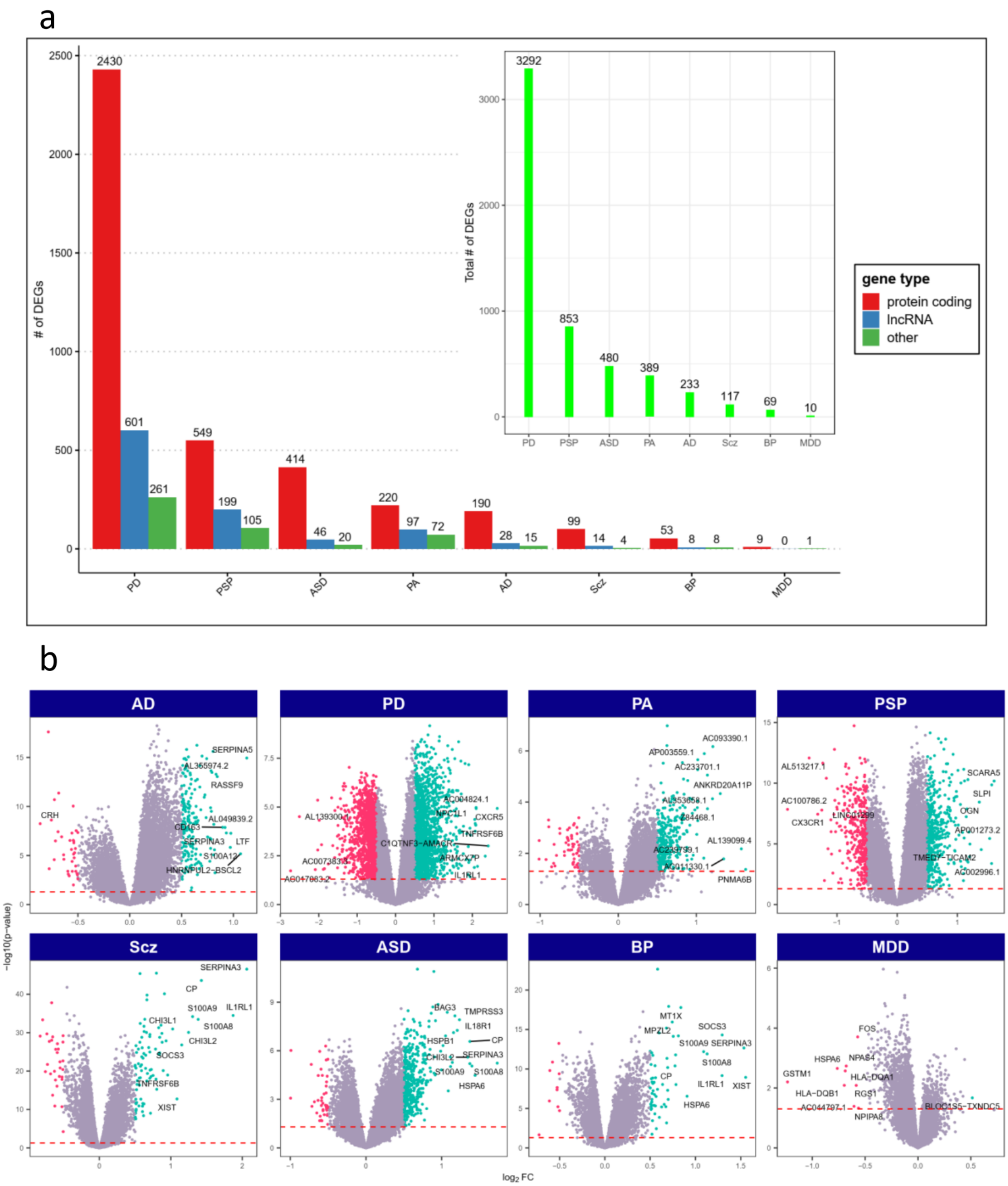


**Figure S11. Condition-specific transcriptome alterations.** (a) The number of differentially expressed genes (DEGs) across conditions. (**b**) volcano plot showing top DEGs labeled for each condition ($P < 0.05$ & |logFC| > 0.5).

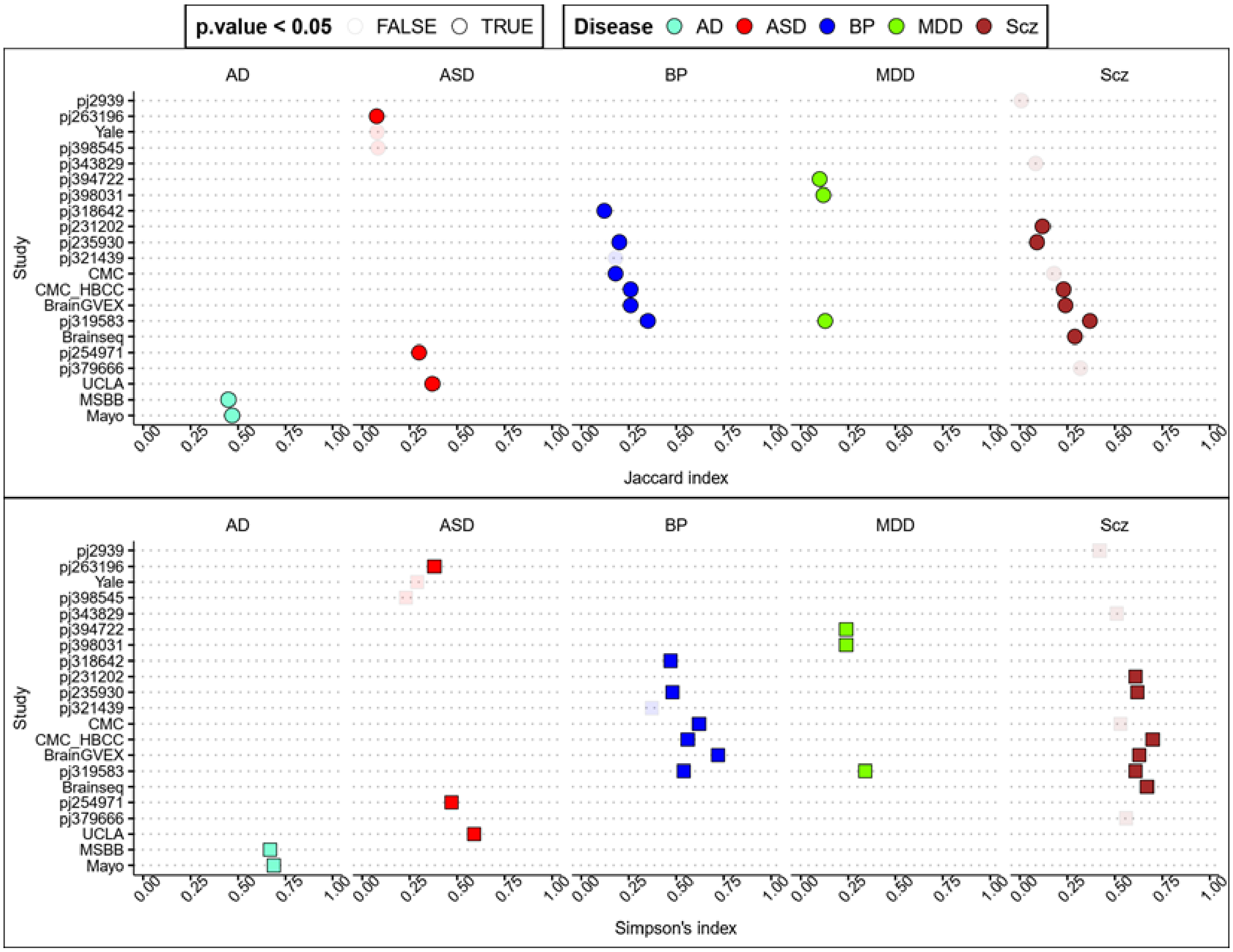


**Figure S12**. **Reproducibility of differential expression results**. Jaccard index (upper plot) and Simpson's index (lower plot) show the similarity ratio of the list of differentially expressed genes for each disease compared to individual datasets.

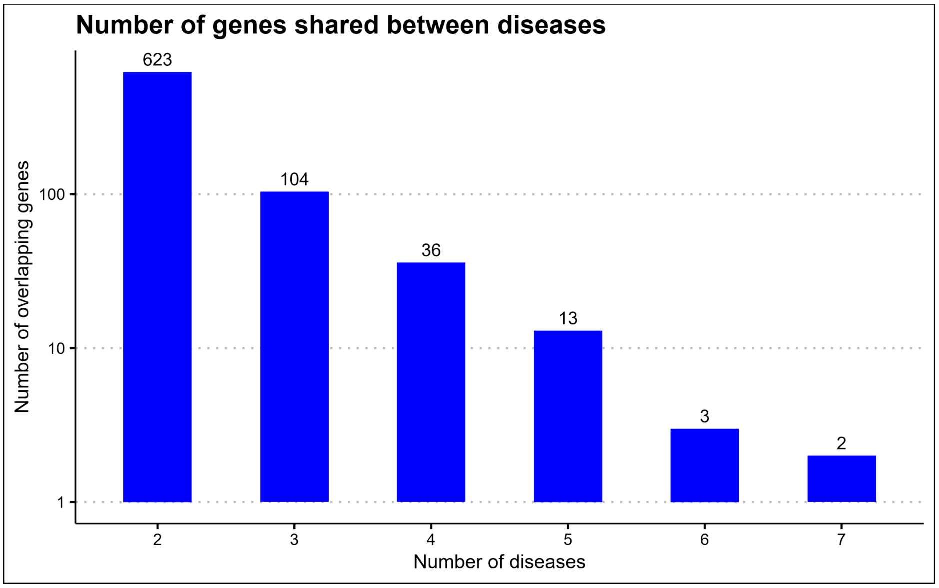


**Figure S13**. The number of differentially expressed genes shared across conditions.

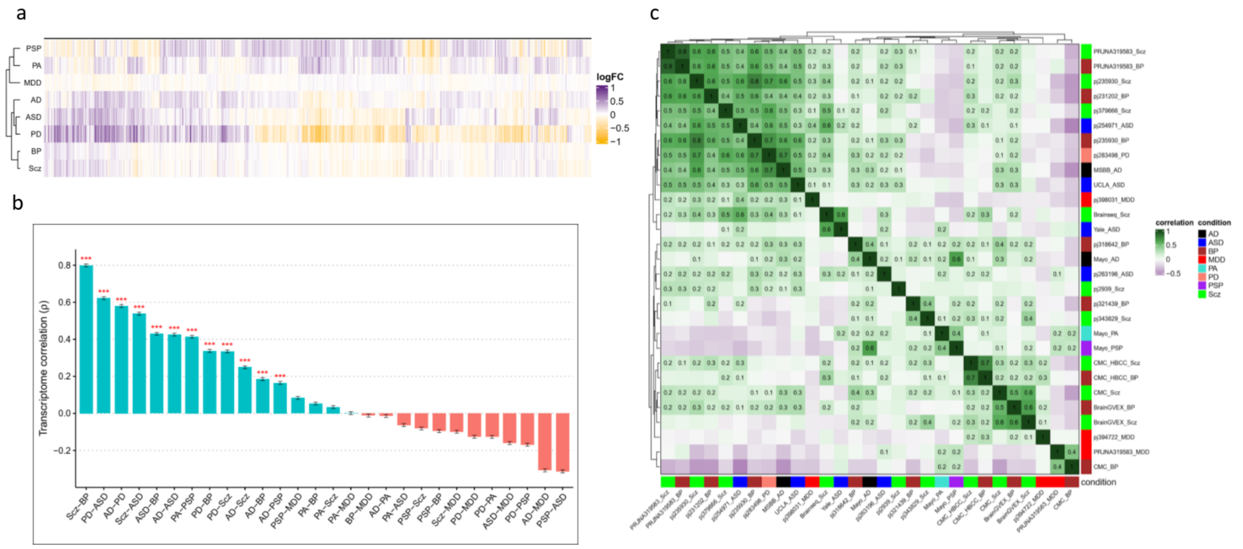


**Figure S14. Transcriptome similarities across eight brain diseases**. **(a)** A heatmap of logFCs from 15819 genes shared between the conditions. (**b**) Barplot shows top pairwise transcriptome correlation across conditions measured by Spearman's correlation of logFC values from common genes (* FDR-corrected P < 0.05). (**c**) Reproducibility of transcriptomic correlations. The heatmap shows Spearman's correlations obtained from the comparison of logFC values of common genes across individual and combined datasets (identified by condition name) for each condition.

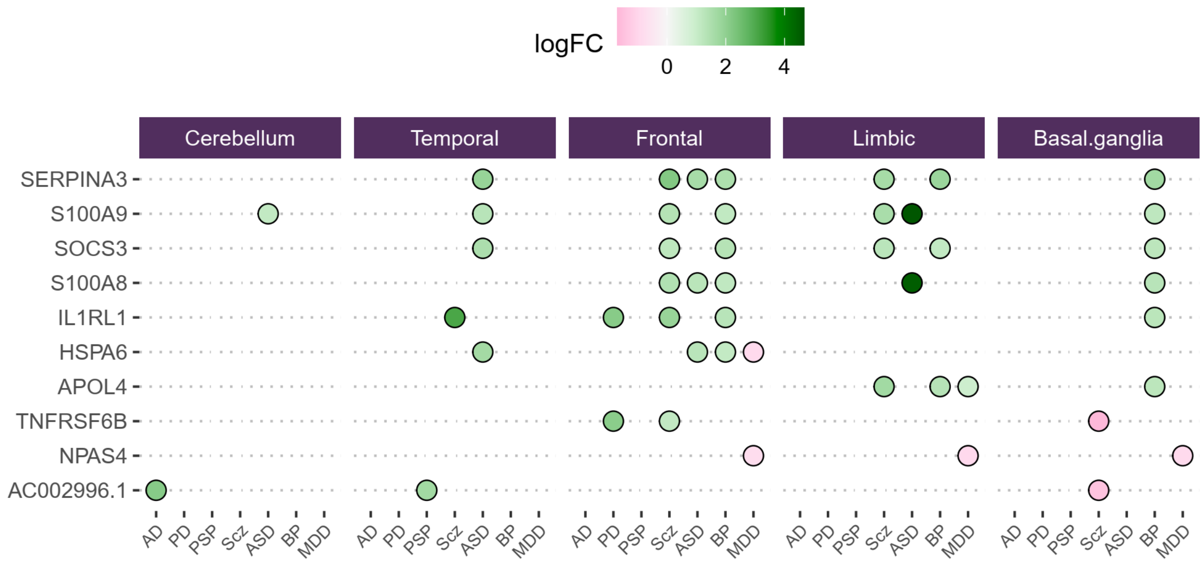


**Figure S15**. Top overlapping genes with differential expression across conditions and brain regions, with color showing logFC value.

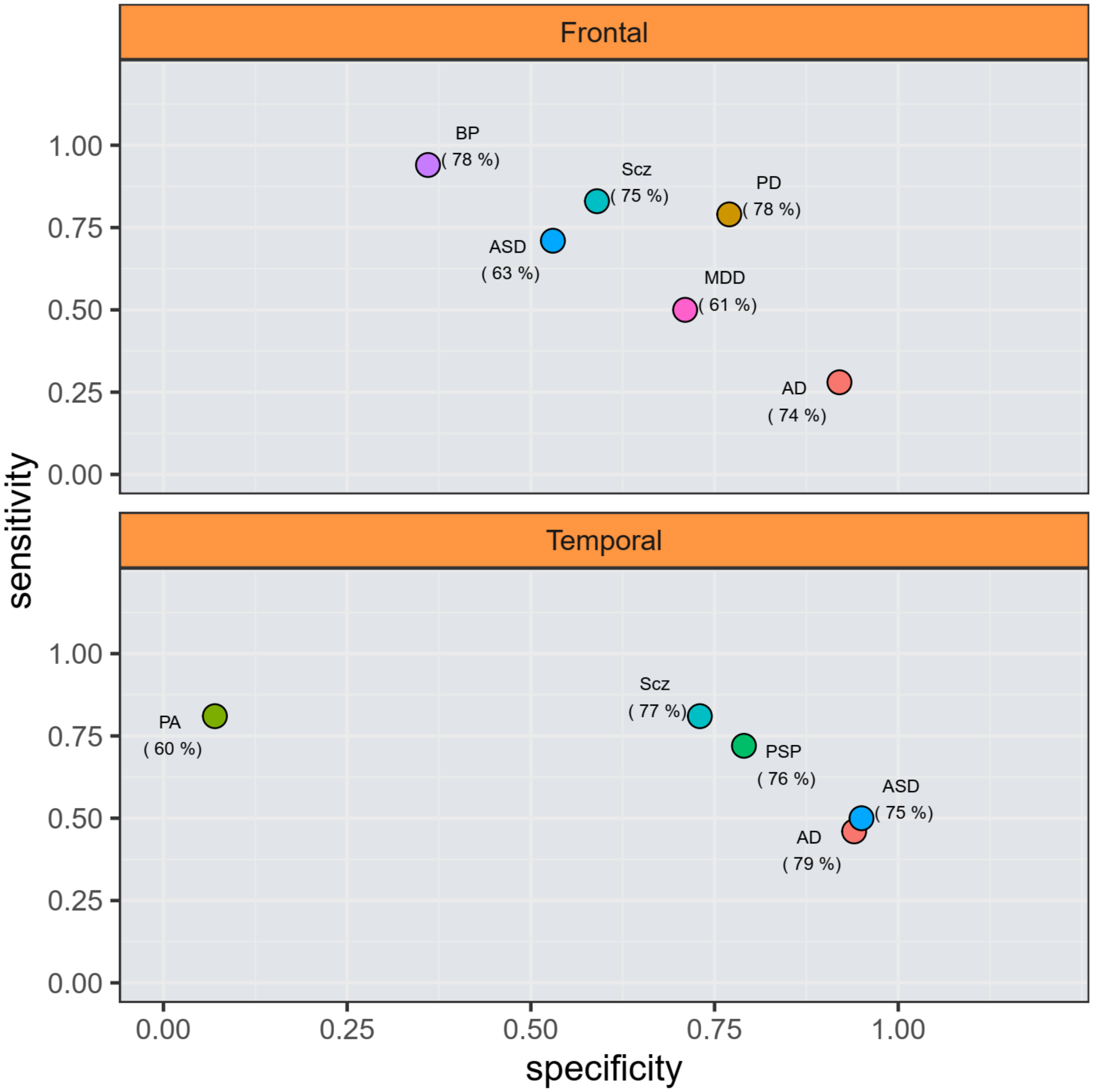


**Figure S16.** Classifier models show the power of frontal and temporal regions to discriminate between transcriptomes of the conditions and control subjects. Each classifier model was built using normalized expression of differentially expressed genes for each region. The *x-axis* shows specificity and the *y-axis* shows the sensitivity of the model. Values in parenthesis show prediction accuracy (%) of each region for each condition.

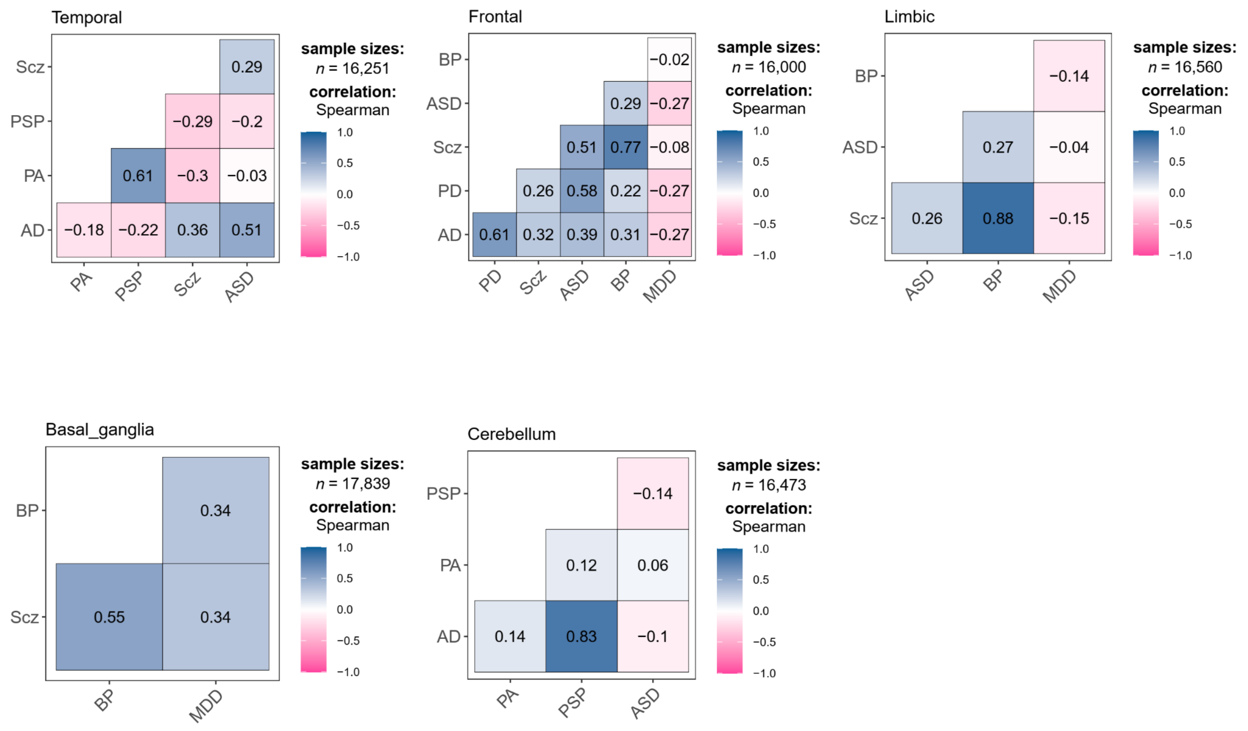


**Figure S17. Brain region transcriptome alterations correlations across conditions.** Transcriptome alterations overlap within each region obtained by Spearman's correlations using logFC values of the shared genes for each region across conditions.

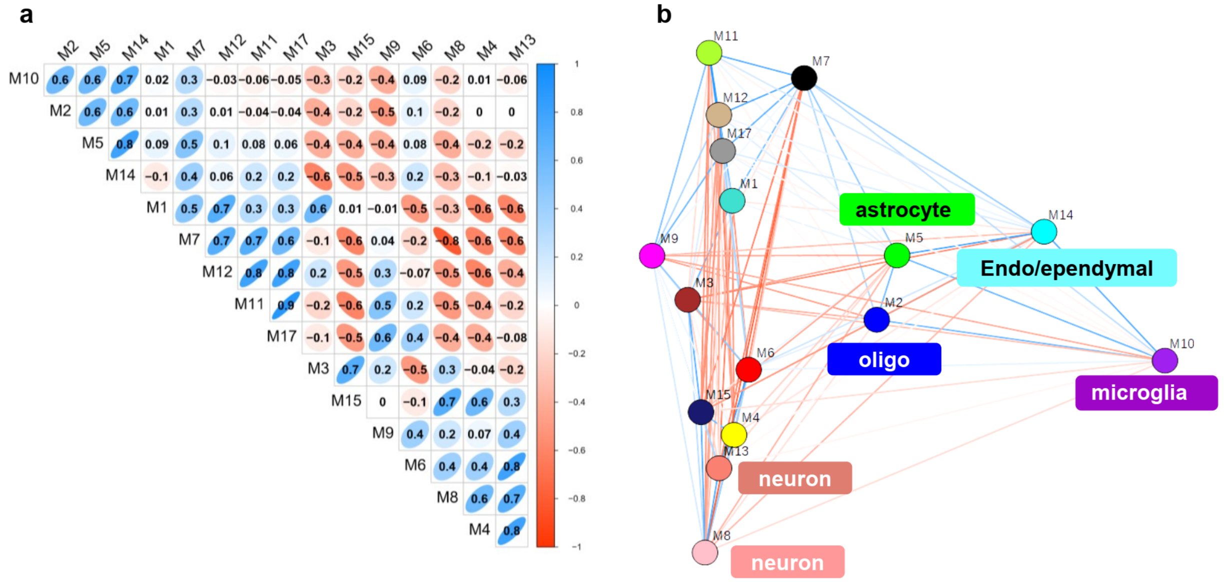


**Figure S18. Co-expression modules relationship.** (**a**) Correlation plot corresponding to obtained from co-expression topological overlap of the modules. (**b**) Multidimensional scaling plot depicts modules relationship, with blue and red color representing positive and negative correlations, respectively.

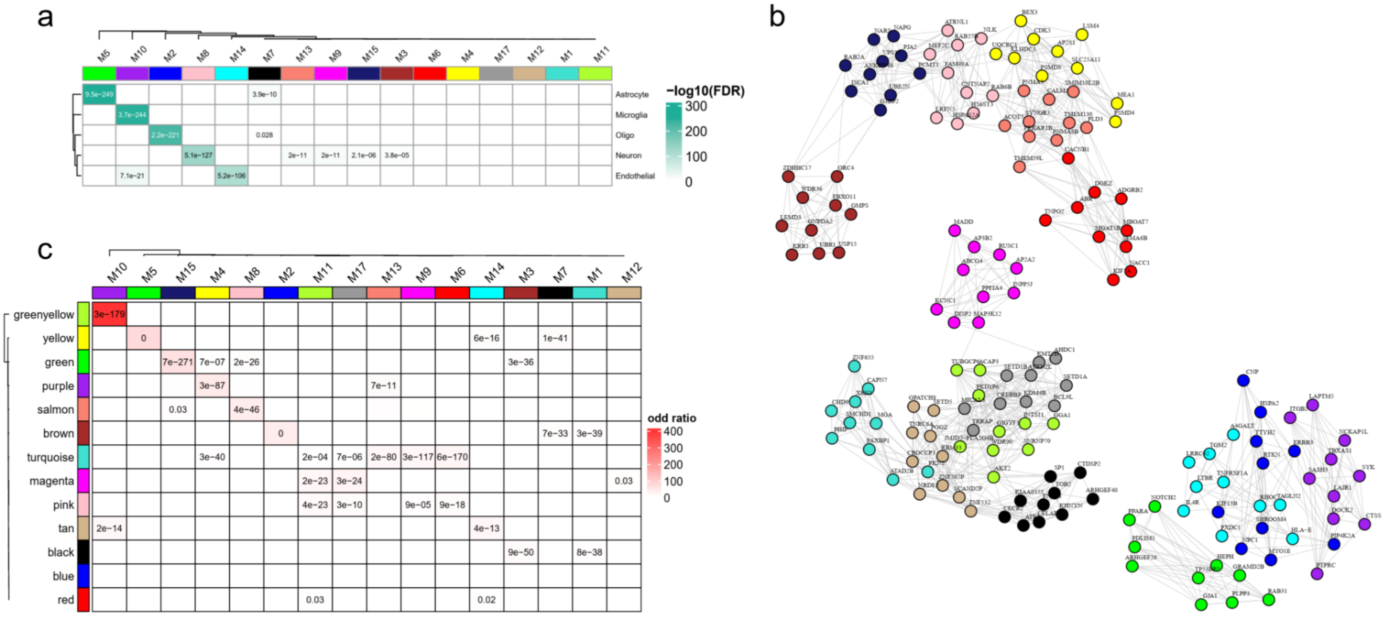


**Figure S19. Co-expression modules characteristics. (a)** Brain cell type-specific enrichment of co-expression modules measured using a single-cell expression dataset composed of five main brain cell types including neurons, astrocytes, oligodendrocytes, microglia, and endothelial cells. Fisher's exact tests were used to perform the comparisons. Values show -$\log_{10}$(FDR-corrected p-values). (**b**) A network of top hub genes (nodes) within each module with highest changes across diseases. Each color shows a module. Edges indicate gene-gene weighted-correlations. **(c)** Overlap of gene modules with network modules from Gandal et al[20]. Color key shows odd ratio and FDR-corrected p-values are shown for significant overlaps (FDR<0.05).

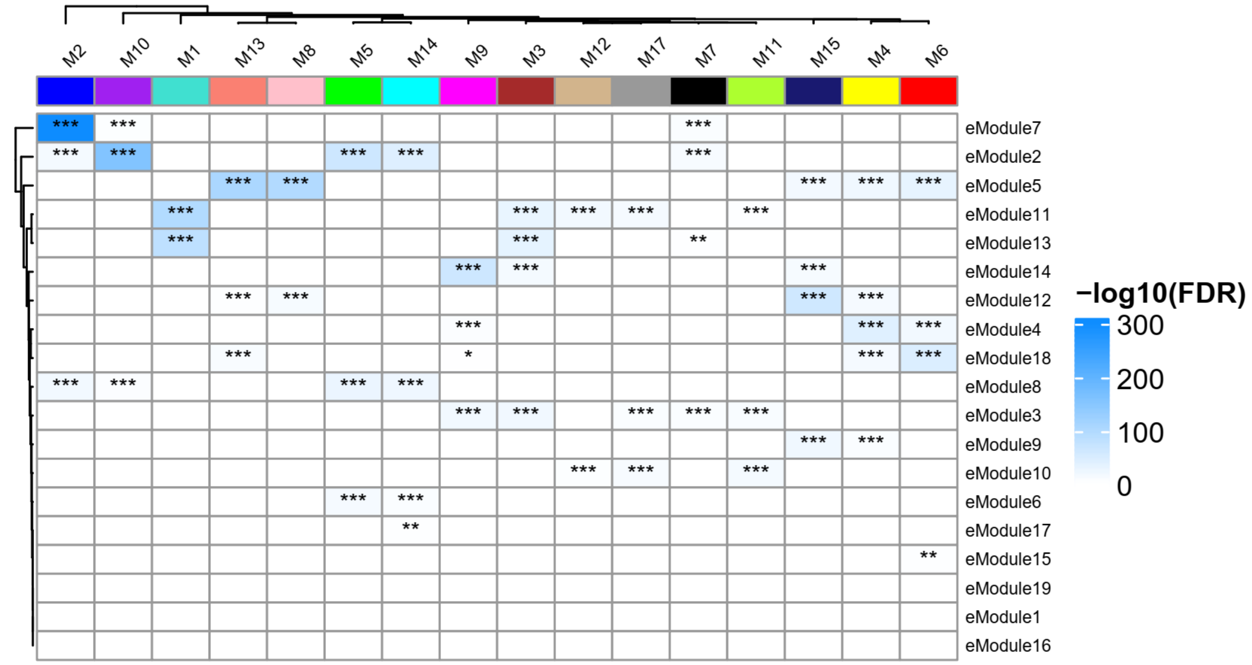


**Figure S20.** Enrichment of co-expression modules for brain enhancer RNAs.

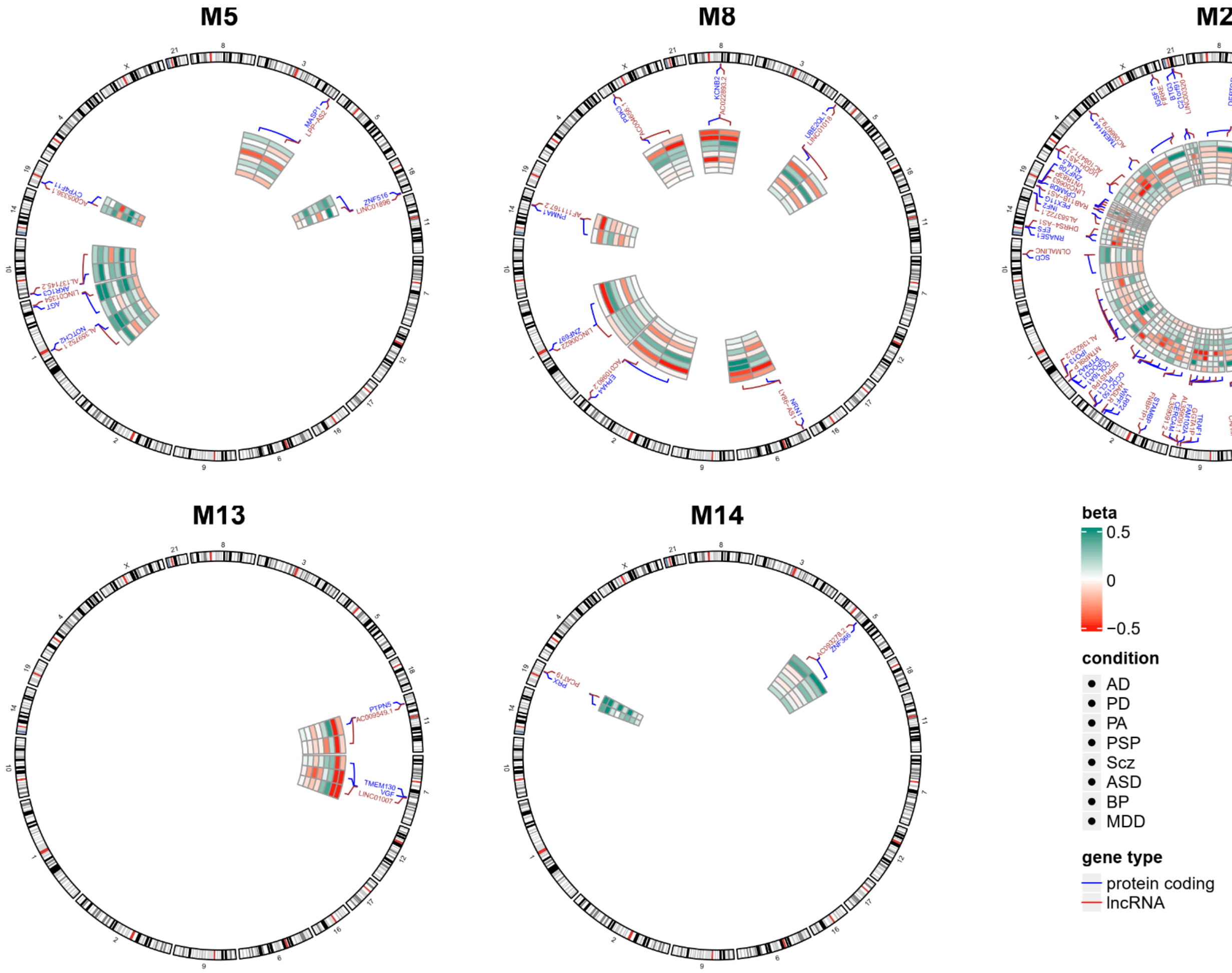


**Figure S21**. A circular heatmap showing expression fold change (beta) of protein-coding genes and their flanking lncRNAs in cell-type-specific modules across conditions.
